# Carbohydrates and the oxidative branch of the pentose phosphate pathway modify *Bacteroides thetaiotaomicron* phage resistance by phase variable S-layers

**DOI:** 10.1101/2025.05.01.651756

**Authors:** Jaime J. Fuentes, Shaleni Singh, Nicholas A. Pudlo, Stacey L. Heaver, Ruth E. Ley, Eric C. Martens

## Abstract

The human gut microbiota consists of hundreds of bacterial species, some of which persist in the presence of lytic phage that infect them. *Bacteroides* employ numerous phase-variable strategies to survive in the presence of phage, including capsular polysaccharides (CPS) and S-layer lipoproteins. We previously reported that a *Bacteroides thetaiotaomicron* strain lacking CPS exhibits almost complete resistance to multiple phages when forced to express the S-layer protein BT1927. However, this strain was only resistant after certain growth conditions, suggesting nutritional variables alter infection and resistance. We grew this strain on various simple sugars and polysaccharides finding that some substrates (fructose, glucose) promote strong resistance to a single phage (ARB25) while others like *N*-acetylgalactosamine (GalNAC) and mucin *O*-glycans increase susceptibility. Mixing fructose and GalNAc indicates the effects of GalNAc are dominant. Despite increasing ARB25 susceptibility, GalNAc did not reduce *BT1927* transcript or protein levels. Instead, GalNAc reduced the amount of BT1927 displayed on the cell surface and increased outer membrane vesiculation. Mutants in any of the 3 steps of the oxidative branch of the pentose phosphate pathway—grown in fructose—behaved similarly to wild-type cells grown in GalNAc, illuminating this pathway in regulation of sugar-mediated phage-resistance. Despite promoting strong resistance, cells grown in glucose/fructose sometimes displayed sub-populations that appeared to completely lack surface BT1927, suggesting another checkpoint exists to control whether this phage defense is deployed. Finally, we show the mucin sugar GalNAc increases susceptibility to several other phage, which has implications for *B. thetaiotaiomicron* persistence in niches like the mucus layer.

**Importance:** The persistence of viruses that infect bacteria (bacteriophages or phages) in the human gut microbiome and their effects on bacterial physiology and host health are active areas of investigation. Our study investigates how various sugars and polysaccharides alter phage susceptibility and resistance in the model gut symbiont *Bacteroides thetaiotaomicron* to lytic phages that are capable of infecting it. Our finding that the mucin sugar, *N*-acetylgalactosamine, and mucin *O*-glycans that contain this sugar reduce *B. thetaiotaomicron* resistance to multiple phages has implications for how this symbiont persists in different gut microhabitats, such as the mucus layer, and which defense mechanisms it can deploy to survive in these niches.

## Introduction

The human gut is home to a community of symbiotic microbes (microbiome) that impart beneficial functions to their host, including digestion and metabolism of nutrients like complex carbohydrates (1, 2), immune stimulation (3–5), and protection against pathogens (6, 7). Individual bacterial species that compose the microbiome often encounter perturbations, including changing diet, severe osmotic stress, immune responses, antibiotics and exposure to bacteriophages (herein, phages) (8–13), all of which could limit the ability of individual bacteria to persist in the community. Phages are the most abundant biological entities in the human gut (14) and target specific species. The presence of phage could have consequences for microbiome composition, as well as the host, if they were able to eradicate their target bacterial species and eliminate its beneficial functions. Further, phage could disrupt community stability by eliminating their target species and indirectly reducing beneficial functions provided by bacteria that rely on the phage-targeted species (15, 16). Despite the potential for phage to cause such disruptions, longitudinal human studies indicate that most bacterial strains are stable for long periods (9, 17, 18). A notable experiment in which phage were administered to gnotobiotic mice colonized with a community of human bacteria demonstrated that although specific bacterial species experienced transient reductions in abundance, their abundance later recovered, and their target phage were eliminated or reduced below detection limits (19). Notably, both bacterial species that survived this phage challenge were members of the genus *Bacteroides*, which is particularly prominent in the microbiomes of people from industrialized countries. This study reveals that at least some gut bacteria have mechanisms to persist in the gut ecosystem despite the presence of phage that prey on them.

Indeed, mechanisms through which gut bacteria resist phage infection and their effect on microbiome stability are emerging. Metagenomic and functional studies have detected a number of known or previously unknown phage resistance mechanisms in gut microbiome species such as: clustered regularly interspaced palindromic repeat (CRISPR) arrays (20–22), cyclic oligonucleotide-based antiphage signaling systems (CBASS) (22–24), restriction-modification systems (22, 25), and variation of cell surface structures like capsular polysaccharides (CPS) and surface layer lipoproteins (S-layers) (22, 26, 27). While mechanistic characterization of these systems is ongoing (20), few have been examined through the lens of regulation in response to the bacterial cell’s nutritional status.

Phase-variation is a reversible phenomenon that often involves DNA inversion of the promoter that directs transcription of a specific gene or operon, such that some cells in a population express the gene and others do not. We previously reported that the model human gut symbiont, *Bacteroides thetaiotaomicron*, is able to resist phage infection through phase-variable expression of CPS biosynthetic loci, candidate S-layer proteins, restriction-modification, and lipoprotein systems (27). Some of these systems have since been reported to be important for phage resistance in other *Bacteroides* species (28, 29). The presence of these phase-variable systems enables *B. thetaiotaomicron* to “pre-adapt” some members of its population to survive phage infection because resistant cells, which already exist in a population (*i.e*., they have turned on one of the mechanisms noted above), survive and become dominant when phages are present. Interestingly, the reversibility of these systems continuously generates susceptible cells that may be killed by phage and potentially sustain the viral population (27), a phenomenon observed decades ago in some *Bacteroides* species and referred to as a “carrier culture” state or pseudolysogeny, although the latter term has been used to refer to different phenomena more recently (30, 31). The presence of reversible phage resistance mechanisms is likely a result of constant coevolutionary adaptation by bacterial species and could result in long-term persistence of phage populations within the gut along with their host bacteria. Consistent with this, phage populations in some human studies have been observed to be stable for at least 1 year (32, 33).

S-layer lipoproteins are ubiquitous among both Gram-positive and Gram-negative bacteria and are characterized by a proteinaceous crystalline-like surface structure (34, 35). Previous work characterized a functional role for one phase-variable *B. thetaiotaomicron* S-layer protein (encoded by the *BT1927* gene), which was visualized as a crystalline-like surface structure by electron microscopy when the *BT1927* gene was expressed (35, 36). *BT1927* transcription is mediated by a phase-variable promoter flanked by invertible repeats that mediate recombination and inversion of the promoter since mutation of one of the two inverted repeats can “lock” the promoter in the ON or OFF position. Antibody staining demonstrated that the BT1927 S-layer gene is expressed in about 1 in 1,000 wild-type cells grown *in vitro* in rich medium (*i.e*., in the absence of phage), while human metagenomic analysis indicated that the promoter-on frequency is higher *in vivo*. While this first study of BT1927 did not test for a role in phage resistance, it reported that BT1927 promotes resistance to complement-mediated killing (36). In our previous study of *B. thetaiotaomicron* phage resistance, increased transcription of *BT1927* in response to phage challenge was a prominent response in the absence of CPS, suggesting it promotes phage resistance. Consistent with this, a strain in which BT1927 was locked on demonstrated nearly complete resistance to at least four phages (27). However, this resistance was only observed when colonies were “aged” for three or more days on solid medium with glucose as the primary carbon source prior to culture in liquid medium. This age-variable resistance occurred even though transcription of the BT1927 gene was locked on, suggesting that growth conditions related to colony aging could either modify the transcriptional output of the *BT1927* promoter or act post-transcriptionally to modify BT1927 effects.

*B. thetaiotaomicron* is adept at harvesting carbohydrates from a variety of diet and host polysaccharides, which are a significant nutrient source for this species compared to protein (37–41). In this study, we measured how the different carbohydrates utilized by *B. thetaiotaomicron* impact the ability of the BT1927 S-layer to promote phage resistance when the *BT1927* gene is constitutively expressed. We found that *B. thetaiotaomicron* exhibits variable resistance to a single lytic phage, when cultured in liquid medium containing one of 27 different individual carbohydrates (simple sugars and polysaccharides). We also discovered that steps in the oxidative branch of the pentose phosphate pathway are important for full phage resistance via BT1927 expression. The carbohydrates that reduced BT1927-mediated resistance most were the sugar *N*-acetylgalactosamine (GalNAc) and mucin *O*-glycans that contain this sugar. Interestingly, decreased phage resistance did not correlate with reduced BT1927 transcription or protein abundance in the cell, but was associated with less BT1927 protein on the cell surface and increased production of outer membrane vesicles (OMVs) that may disrupt the integrity of this protective S-layer. Because the carbohydrate-based nutrients available to gut bacteria frequently change with variations in host diet, thereby influencing their cellular responses, our findings have important implications for understanding how the stability and co-existence of bacteria and phage populations in the human gut may be influenced by diet or bacterial occupation of niches like the colonic mucus layer.

## Results

### Carbohydrates influence BT1927 S-layer mediated phage resistance

The nutritional environment experienced by a bacterial cell, such as the presence of specific nutrients or nutrient deprivation, has potential consequences for interactions with phage. Two classic examples of bacterial nutrient exposure that modify phage susceptibility in *E. coli* are the presence of maltose or iron-deficient conditions, each of which induces expression of a cognate phage receptor (LamB or FhuA, respectively) (42, 43). Given the previously determined requirement for “aging” BT1927-expressing cells for three days to elicit strong protection against ARB25 phage, we hypothesized that an unknown nutrient signal(s) alters the ability of BT1927 to protect against phage. Consistent with our previous observations (27), when a strain with the phase-variable *BT1927* promoter genetically locked-on (herein, BT1927:ON) was aged for 3 days and infected with ARB25, it exhibited similar growth as a heat-killed phage control when infections were performed in media containing glucose (**Figure 1A**). This protection against ARB25 was dependent on both locked expression of BT1927 and colony aging, since both a BT1927 locked-off strain (BT1927:OFF) and the BT1927:ON strain grown without aging achieved significantly lower growth (P<0.0001) at 24 hours and generally failed to maintain the same high density as the BT1927:ON aged strain in the first ∼40 hours of infection (**Figure 1A, B**, compare red and blue curves to green). Growth eventually increases later after infection with many strains or conditions, likely driven by the emergence of cells expressing other phase-variable resistance proteins, of which there are 7 known to exist in the engineered strain used here.

**Figure 1.**
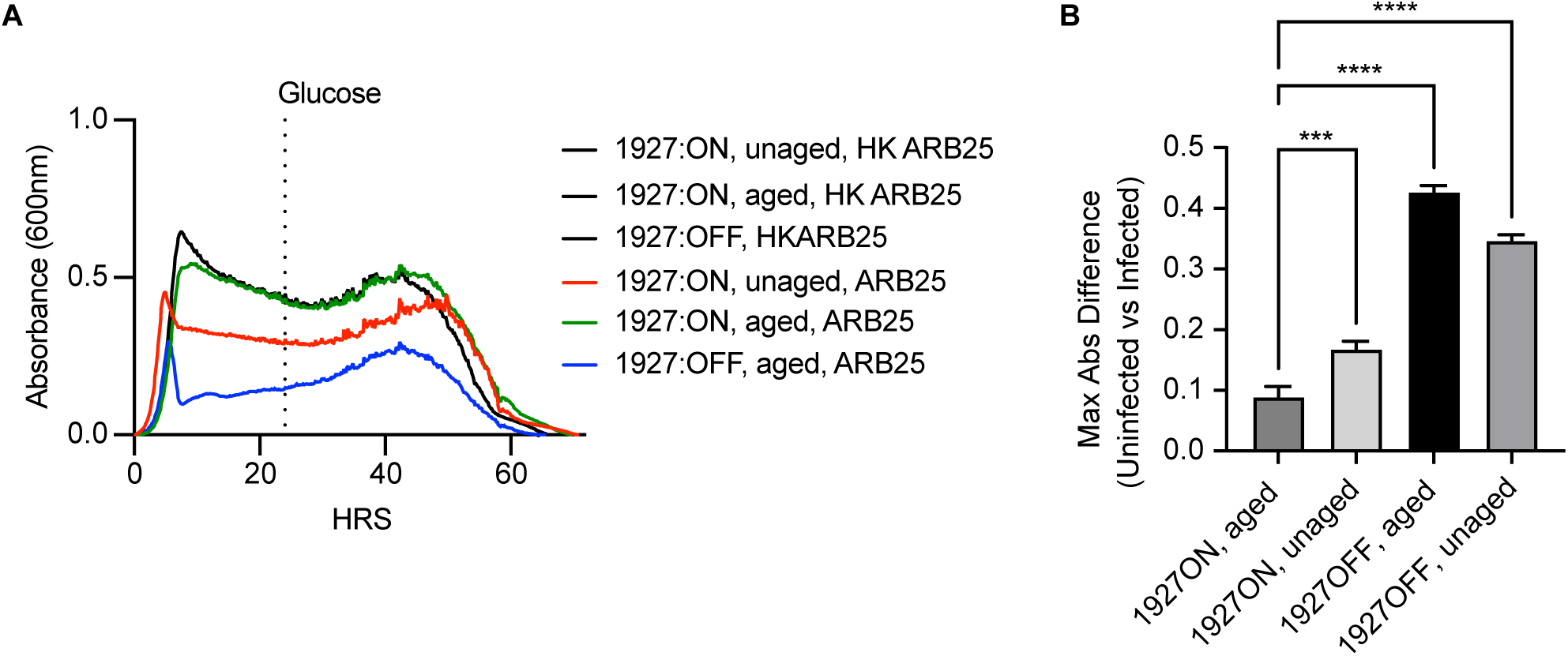
Colony aging increases BT1927-mediated resistance to ARB25 phage. (**A**) Growth of unaged (red), aged (green) BT1927:ON strain or BT1927:OFF (blue) grown in media containing glucose and challenged with viable ARB25 or heat-killed (HK) ARB25 (black curves). (**B**) Maximum absorbance differences of HK ARB25 (uninfected) versus ARB25 challenged unaged or aged BT1927:ON or BT1927:OFF strain, calculated at 24-hours (One way ANOVA, **** P<0.0001, *** P=0.0009).

Consistent with previous work (36), scanning electron microscopy (SEM) imaging of the BT1927:OFF strain did not reveal a lattice structure on the outside of the cell (**Figure 2A**). However, individual BT1927:ON cells did display a surface lattice that is similar to what has been previously observed (**Figure 2B).** Interestingly, images of a phage-free culture of the BT1927:ON strain grown in medium with glucose as the main carbohydrate source contained a population of ∼60% of cells that appeared to display the lattice structure on the majority of the cell surface while the remaining individual cells appeared mostly devoid of this structure, suggesting that there may be variation in the expression of the BT1927 S-layer at the single cell level (**Figure 2C, Figure S1A**).

**Figure 2.**
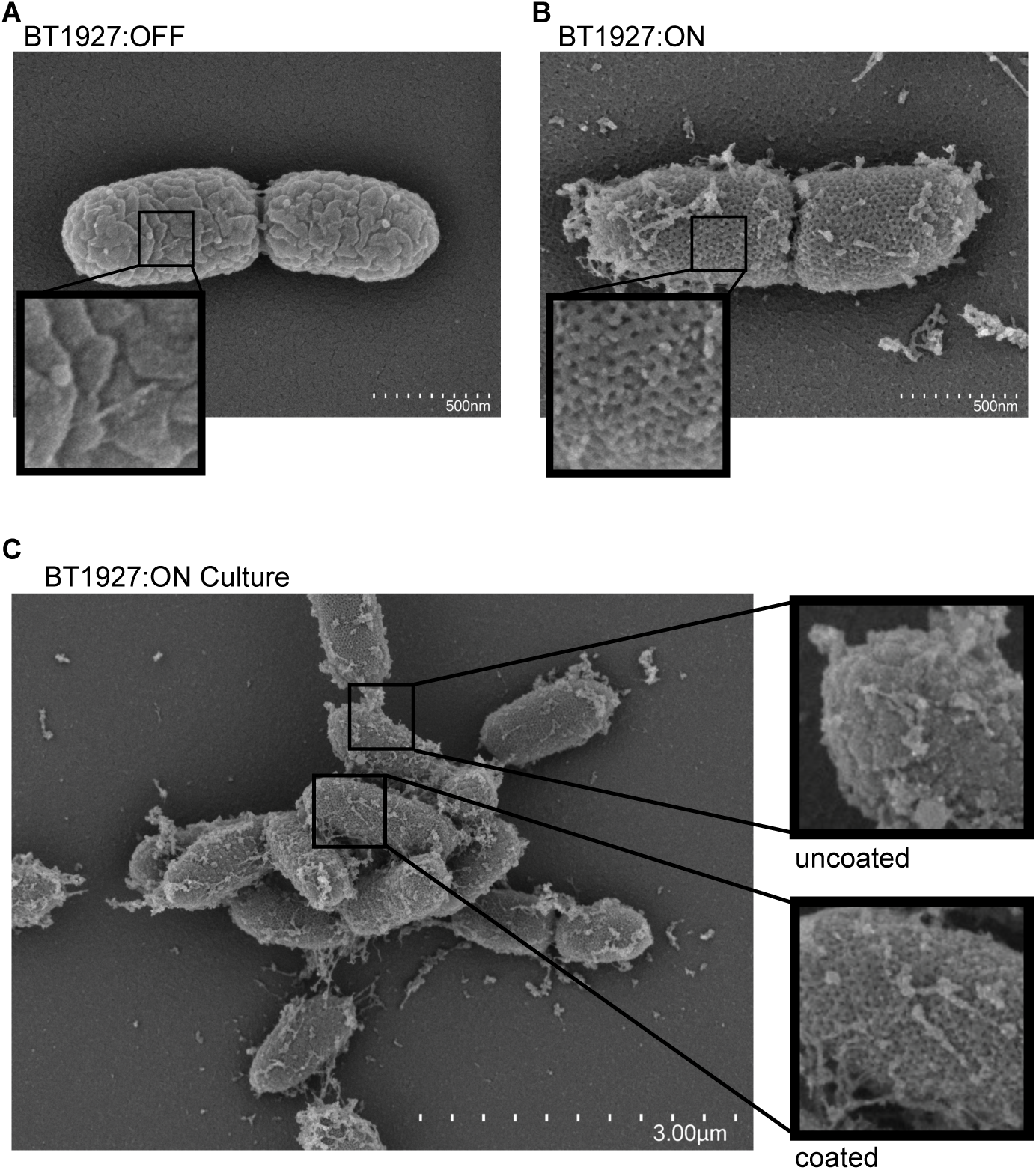
BT1927 expression is associated with a crystalline surface structure. SEM images of the BT1927:OFF strain (**A**) or BT1927:ON strain (**B**). (**C**) Multiple cells of the BT1927:ON strain showing the presence of both coated and uncoated cells. Enlarged inlaid images are provided for each panel to highlight presence or absence of structural features and uncoated or coated cells.

To explore how the aged colony, high ARB25-resistance phenotype persists over time with repeated liquid culture, aged colonies were grown in liquid BPRM glucose, allowed to grow for 24 hours, and sub-cultured 1:100 each day for 5 days. Each day, a portion of the culture grown in BPRM glucose for the previous 24 hours was also sub-cultured and subjected to ARB25 infection (MOI 0.5). Interestingly, we found that passaging of cultures for two days or longer in liquid BPRM-glucose medium was sufficient to promote strong resistance even when starting with cultures of BT1927:ON that had not been grown on solid medium (**Figure 3A, Figure S1B, C**). This observation suggests that multiple generations of growth in glucose—and not necessarily aging on plates—is sufficient to increase the effect of BT1927 to protect against ARB25. Given that glucose is used as the sole carbohydrate source in the media that promoted this phenotype, we aimed to determine how other dietary carbohydrates might alter BT1927-mediated resistance as carbohydrates are the major group of nutrients used by *B. thetaiotaomicron*. We leveraged a panel of 14 monosaccharides and 13 polysaccharides that support the growth of *B. thetaiotaomicron* when present as sole carbon sources and measured growth dynamics of the BT1927:ON strain during infection with ARB25 phage. Briefly, the BT1927:ON strain was grown on BPRM solid agar medium containing glucose for three days, colonies were cultured in liquid BPRM glucose, allowed to grow for ∼20 hours, and sub-cultured into BPRM containing each carbohydrate followed by phage challenge (**Figure S1D**).

**Figure 3.**
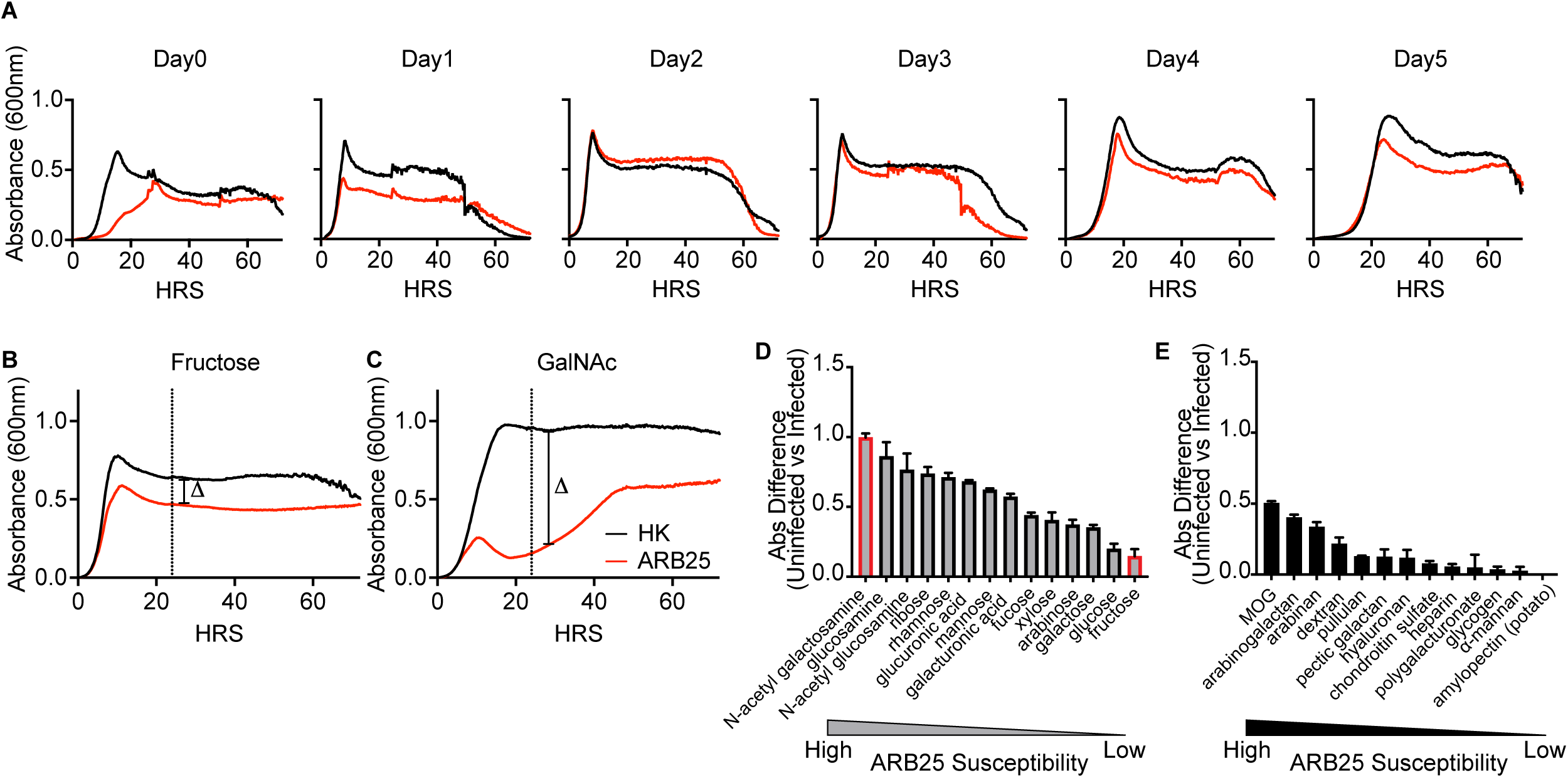
Different carbohydrates modify susceptibility to ARB25 phage. (**A**) Growth of the unaged BT1927:ON strain treated with viable ARB25 (red) or HK ARB25 (black), passaged for 5 days in glucose. (**B**) Growth of the BT1927:ON strain in media containing Fructose (**C**) Growth of the BT1927:ON strain in media containing GalNAc. The delta symbol in C and D indicates the absorbance difference measured to quantify growth differences between ARB25 and HK. (**D**) Ranked monosaccharide results in order of higher to lower ARB25 susceptibility. (**E**) Ranked polysaccharide results in order of higher to lower ARB25 susceptibility.

Several of the carbohydrates in our panel resulted in growth in the presence of ARB25 that was similar to glucose. For example, fructose also promoted a strong and uninterrupted early growth increase in the presence of ARB25 (**Figure 3B, Figure S2A**). In contrast, other carbohydrates elicited growth in the presence of ARB25, that was more similar to the unaged or BT1927:OFF strain grown in glucose (**Figure 1A**). For example, growth in medium containing *N*-acetylgalactosamine (GalNAc) led to an initial growth increase after ARB25 exposure that was blunted and remained low for the first ∼40 hours after initial infection (**Figure 3C**). This low early growth on GalNAc suggests that, in contrast to glucose and fructose, GalNAc fails to promote and perhaps even suppresses BT1927-mediated resistance. As observed with unaged or unpassaged colonies, cultures in GalNAc eventually gained in absorbance (**Figure 3C**), likely due to the emergence of cells that express other resistance mechanisms. Our previous study demonstrated that up to 10 different phase-variable systems (excluding polysaccharide capsules) could be expressed *in vitro* in the presence of ARB25 challenge (27): 8 putative S-layers (including BT1927), a restriction-modification system and a single locus encoding 3 lipoproteins different from the S-layers. Therefore, we probed transcription of these nine other phase-variable systems, plus locked on *BT1927*, during growth in GalNAc and fructose and in the presence of ARB25. This experiment revealed that fructose promoted the emergence of cells with increased transcription of the genes encoding the seven other S-layers, the restriction-modification system (*BT4013-15*) and the additional lipoprotein locus (*BT0291-94*). This suggests that these different systems may confer additional or better protection beyond BT1927 and are therefore selected later in culture in the presence of ARB25. Interestingly, five of the nine systems expressed in fructose had significantly lower expression in GalNAc, and one gene encoding a candidate S-layer (BT1507) was expressed at significantly higher levels in GalNAc (**Figure S2B, C**). These results support the conclusion that different sugars modify the range of phage resistance responses that emerge in a population, possibly by modifying the effectiveness of each specific mechanism during growth in a given sugar.

Since we observed a range of responses to ARB25 during early infection on different carbohydrates, we quantified the difference between heat-killed and ARB25 infected cultures using 24-hours as a common “early” time point (delta symbols in **Figures 3B, C** indicate examples of the growth difference measured for each carbohydrate). Ranking the effects of individual carbohydrates revealed a continuum of effects on BT1927-mediated resistance, with the previously mentioned fructose and GalNAc exhibiting the highest and lowest resistance, respectively (**Figure 3D**). The effects of monosaccharides tended to be stronger than polysaccharides at reducing BT1927-mediated resistance, although both groups showed a broad range of responses. The polysaccharide that resulted in the most reduced resistance to ARB25 was mucin *O*-glycans (MOG), a mixture of oligosaccharides that interestingly contains the monosaccharide GalNAc (**Figure 3E**). Other monosaccharides present in MOG (galactose, fucose, and *N*-acetylglucosamine) did not increase susceptibility to ARB25 to the same level as GalNAc, although the latter two sugars elicited strong susceptibility in some experiments (**Figure S1D**). There was also discordance between the effects of glucose and the four polymers that contain glucose (dextran, pullulan, glycogen, and amylopectin). Thus, we cannot conclude that polysaccharides universally exert the same effect as the monosaccharides they contain. Since we determined that MOG and GalNAc reduce resistance to ARB25, we tested 5 other phages, finding that growth in these two carbohydrates also reduced resistance to 4 of the phages tested when cultured similarly to the experiments described above with ARB25 (**Figure S2D**).

Given the disparate effects of fructose and GalNAc on resistance to ARB25, we aimed to determine if the strong resistance developed after passage in a medium containing glucose could be reversed by the addition of carbohydrates like GalNAc and if one sugar-associated phenotype is dominant over the other. Passage of cultures in glucose for five days, which promoted strong ARB25 resistance, followed by a shift to GalNAc, decreased growth in the presence of ARB25 (**Figure S3A**), demonstrating that GalNAc can induce increased ARB25 susceptibility in cultures that previously exhibited resistance and consequently suggesting that the resistance is not permanent as would be expected for genetic mutation in the ARB25 receptor or other function required for infection. Continuous passage in GalNAc for 5 days resulted in slightly more blunted growth, suggesting a potential compounding effect. In contrast, passage in fructose resulted in continued strong resistance as indicated by lower difference in maximum absorbance between infected and uninfected bacteria (**Figures 4A, S3A,** note that the resistance at 24h for fructose in this experiment may be underrepresented at some days due to some of the uninfected conditions increasing absorbance on this sugar after the initial entry into stationary phase).

**Figure 4.**
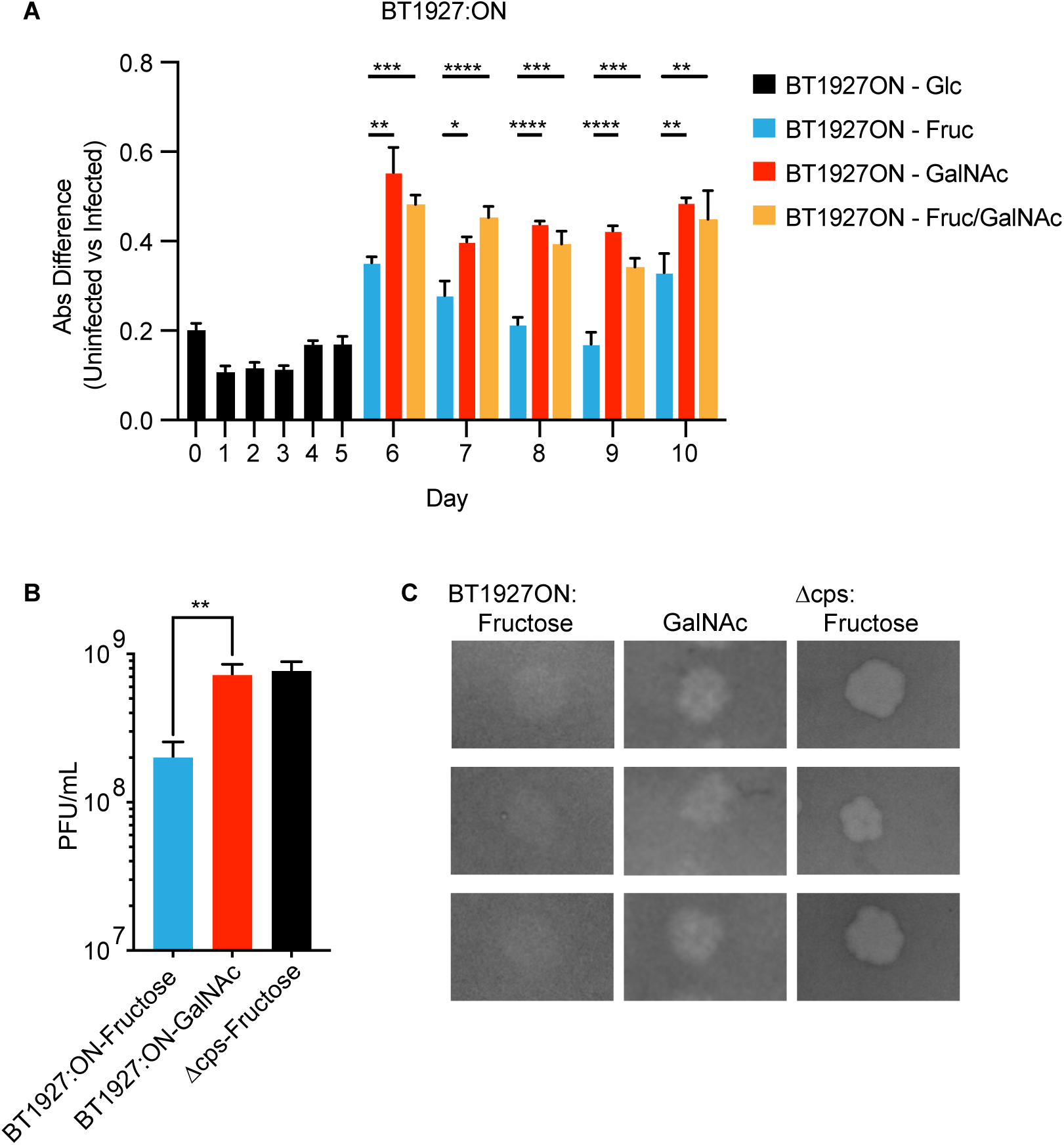
GalNAc increases susceptibility to ARB25 phage. (**A**) Maximum absorbance differences from passaged growth experiments for 5-days in medium containing glucose (black), fructose (blue), GalNAc (red), or a 1:1 mixture of fructose and GalNAc (orange) at 5mg/ml each, n = 3 per condition (One way ANOVA, **** P<0.0001, *** P=0.0002, ** P<0.005, *P<0.05). (**B**) Quantification of plaque forming units with indicated strains grown in top agar containing fructose (blue) or GalNAc (red) compared to the more susceptible acapsular control (Δcps; black), n = 3 per condition, (one-tailed t-test, **P=0.0066). (**C**) Representative images of plaques formed under the indicated conditions and strains as depicted in B.

To address whether fructose or GalNAc are dominant, we passaged cultures in medium containing equal amounts of fructose and GalNAc (5mg/ml each). This resulted in growth that was very similar to GalNAc grown cultures, suggesting that the effect of GalNAc is dominant over that of fructose (**Figure 4A, Figure S3A**). We next determined whether growth on solid medium with GalNAc for 3 days resulted in increased ARB25 susceptibility after cells were grown in liquid medium and exposed to phage. Consistent with our results in liquid medium, cultures from colonies picked from agar containing glucose, first cultured in BPRM glucose, and then infected in BPRM GalNAc showed lower resistance at 24h but later increased growth (**Figure S3B**). In contrast, growing BT1927:ON strain on GalNAc agar plates and subsequently in liquid BPRM GalNAc for growth and infection resulted in cultures that were more susceptible at 24h and failed to increase growth as much as glucose grown cells at later times (**Figure S3C**). This latter result supports our conclusions from repeated culture in liquid medium that previous exposure to glucose exerts an effect that persists for some amount of time after first exposure to GalNAc and this influences how the cells resist ARB25 infection.

We next measured if growth in medium containing fructose or GalNAc alters the plaquing efficiency of ARB25 on the BT1927:ON strain, which could explain the apparently high and low ARB25 resistance trends we observed in liquid cultures. Consistent with our results in liquid media, plaquing efficiency was significantly higher in GalNAc-containing plates compared to those containing fructose and the GalNAc medium promoted a similar number of plaques on the BT1927:ON strain as the parent acapsular strain—which does not have BT1927 locked on—grown on fructose (**Figure 4B**). Moreover, plaques appeared less turbid in the GalNAc medium compared to glucose, potentially indicating that this sugar increases the efficiency of ARB25 infection of individual cells as waves of post-infection phage bursts encounter new prey in their surroundings (**Figure 4C**). Similar to earlier studies showing that the BT1927 locked-on strain is protected against complement killing compared to the locked-off strain in medium containing glucose (36), cultures grown in GalNAc exhibited significantly reduced survival compared to fructose grown cells when exposed to complement (**Figure S3D**). These data further suggests that the protective nature of the BT1927 S-layer is influenced by the specific carbohydrates encountered by the cell.

### N-acetylgalactosamine reduces BT1927 expression on the bacterial surface

We next determined if cultures of BT1927:ON grown in fructose or GalNAc exhibit variations in surface exposed BT1927. We used flow cytometry to measure surface expression of a previously published BT1927 variant with a FLAG epitope appended to the C-terminus (36) (BT1927:ON^F^). To first determine if the presence of the FLAG epitope disrupts BT1927 function with respect to surviving phage, we challenged cultures of the BT1927:ON^F^ strain with ARB25 and determined that it maintains resistance when cultured in fructose (**Figure S3E)**. We then performed flow cytometry using the BT1927:ON^F^ strain grown in fructose or GalNAc to mid-log phase. Using an antibody against the FLAG epitope, the BT1927:ON^F^ strain grown in fructose could be clearly distinguished from the BT1927:OFF strain grown in the same medium by flow cytometry (**Figure 5A**). The BT1927:ON^F^ strain grown in GalNAc was clearly different from fructose-grown cultures and had less intense staining overall with an average geometric mean fluorescence intensity of 200±62 (GalNAc) versus 578±64 (fructose; P = 0.0017, one tailed t-test), although it was also clearly increased relative to the BT1927:OFF control (**Figure 5A)**. This result suggests that GalNAc-grown cells express less BT1927 on the cell surface.

**Figure 5.**
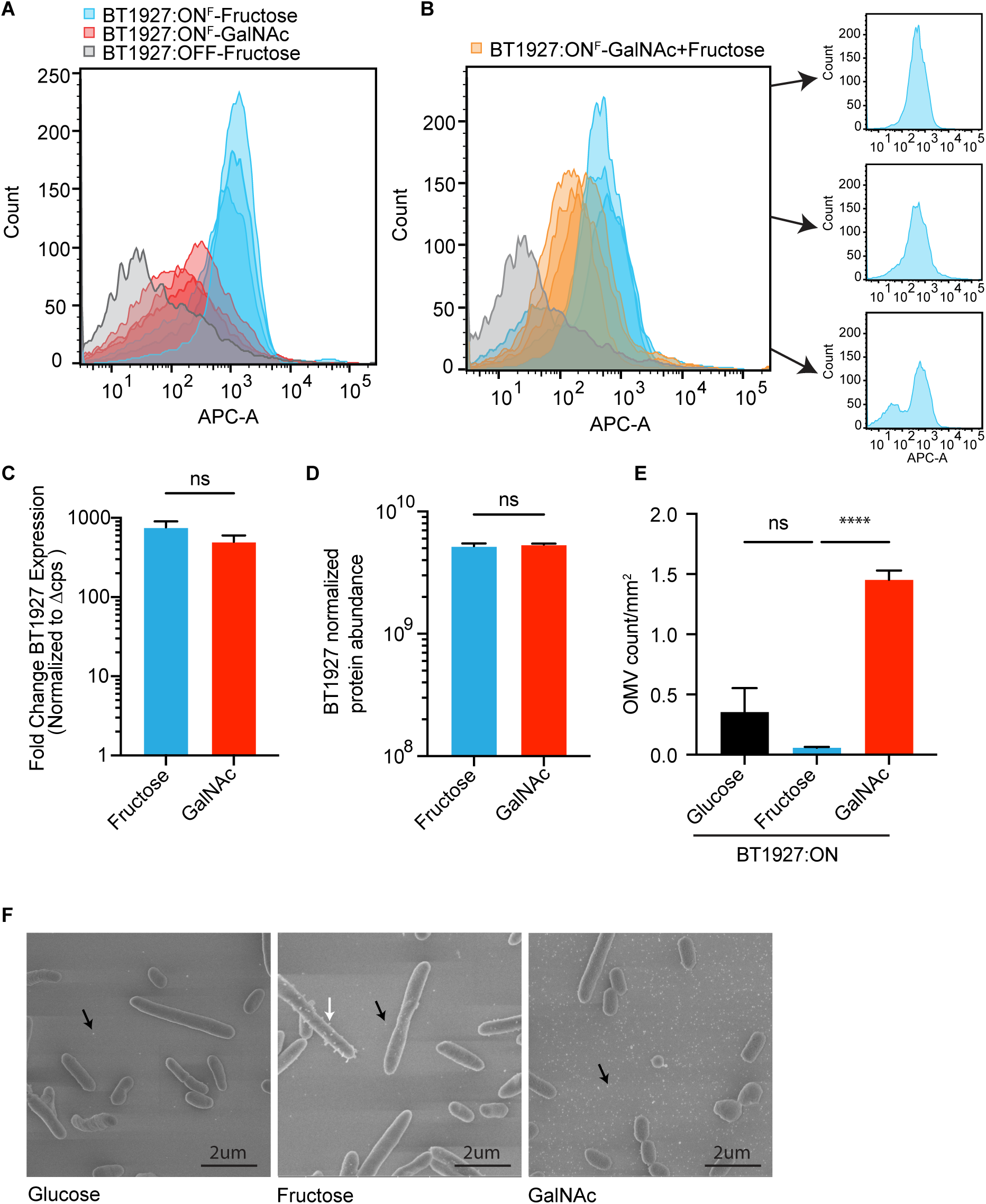
BT1927 surface protein abundance is modified by GalNAc. (**A**) Flow cytometry histograms for BT1927 expression indicated by intensity of FLAG epitope staining (APC-A) in the BT1927:OFF^F^ (grey) strain grown in fructose or the BT1927:ON^F^ strain grown in medium with either fructose (blue) or GalNAc (red). (**B**) Same conditions as depicted in A but with the addition of staining of cells grown in a mixture of GalNAc and fructose (5mg/ml each; orange) instead of GalNAc only. The smaller plots at right show the three individual plots of growth in fructose in which cell population (bottom) displayed a sub-population of cells that stain similarly to the BT1927:OFF^F^ strain. (**C**) BT1927 transcript expression in the BT1927:ON strain grown in either fructose (blue) or GalNac (red). (**D**) Normalized protein abundance in BT1927:ON strain grown in fructose (blue) or GalNac (red). (**E**) OMV particle count per mm^2^ from cells grown in either glucose (black), fructose (blue) or GalNAc (red) (Welch ANOVA, **** P<0.0001). (**F**) representative images for each condition in E. Arrows indicate apparent OMV particles either cell associated (white) or free from cells (black).

An interesting observation in several repetitions of this experiment, was the presence of a distinct population of BT1927:ON^F^ cells that stained similarly to the BT1927:OFF control population after cultivation in fructose (**Figure 5B** small histograms, **Figure S4A**). In the first experiment in which this occurred (**Figure S4A**), all three biological replicates were similar to each other while in our first experiment all three replicates lacked this population (**Figure 5A**). We interpret this observation to indicate that in some cases, a population of cells can be present that expresses very little cell-surface associated BT1927 despite the fact that expression is locked on and the growth sugar (fructose) promotes overall strong ARB25 resistance in the population. We do not suspect that this is a technical artifact (*e.g*., poor antibody labeling in the experiment with a prominent BT1927 low population) because the BT1927 high populations displayed similarly high staining intensity. Instead, this bimodal population of BT1927-expressing cells could reflect the two populations of cells that we initially observed by SEM imaging of cells grown in glucose (**Figure 2C**). If true, these two observations would support the conclusion that *B. thetaiotaomicron* possesses a switch or decision-making mechanism that governs the fate of individual cells to express BT1927 (and perhaps other proteins) on the cell surface. To further investigate variation in BT1927 expression between experiments with growth in fructose, we grew the BT1927:ON^F^ strain in several different anaerobic chambers with each replicate cultivated from a different starting colony. Among the four experiments, one exhibited a low staining population of BT1927:ON (**Figure S4B**), supporting the conclusion that this is a repeatable phenomenon that could be driven by an unknown environmental stimulus or stochastic variation in *B. thetaiotaomicron* cell populations. Since our previous data indicated that GalNAc was dominant in decreasing the resistance of the BT1927:ON strain to ARB25 when fructose was also present, we grew the BT1927:ON^F^ strain in media containing both fructose and GalNAc to measure BT1927 expression levels. Strikingly, the BT1927:ON^F^ strain grown in a mixture of fructose and GalNAc was clearly distinct from fructose-grown cultures, had less intense staining overall and was very similar to cells grown only in GalNAc (**Figure 5B**). This observation suggests that the effect of GalNAc is dominant over fructose and may not allow maximum expression, asembly or retention of BT1927 on the cell surface.

Since GalNAc reduces the amount of surface associated BT1927 protein, we next sought to determine at which point in expression this effect is exerted. Since the *BT1927* promoter is already locked in the phase-variable “on” position, we measured if transcription of the *BT1927* S-layer is downregulated upon exposure to GalNAc. Comparison of the BT1927:ON strain in GalNAc or fructose (in the absence of phage) revealed that *BT1927* transcript was not diminished by GalNAc (**Figure 5C**). Similarly, high *BT1927* transcription was observed during growth on another carbohydrate, glucosamine, that increases ARB25 susceptibility (**Figure S4C**) Next, we performed a western blot on the BT1927:ON^F^ strain grown in media containing either fructose or GalNAc to assess BT1927 protein levels in whole cells. BT1927 protein did not appear to be different between fructose and GalNAc grown cultures (**Figure S4D**), although the western blot was not performed quantitatively and is unlikely to discriminate between small changes in protein expression that might explain the partial reduction in surface staining observed by flow cytometry. To more quantitatively determine if these two sugars alter BT1927 protein abundance, we performed proteomics using LC-MS/MS to measure the relative normalized protein abundance of BT1927 in the BT1927:ON strain grown in either fructose or GalNAc. The abundance of BT1927 was not significantly decreased by growth in GalNac compared to fructose (**Figure 5D)**. As a control for expected sugar-specific responses, growth in fructose led to a 1.6-5.0-fold increase in five of the eight individual proteins encoded in a previously identified polysaccharide utilization locus (PUL) for the fiber levan that is upregulated in response to fructose (44) (**Figure S4E**). Taken together, although GalNAc increases ARB25 susceptibility and decreases its abundance on the cell surface, we do not have any evidence that GalNAc reduces transcription or translation of the BT1927 lipoprotein, which therefore necessitates that lack of surface expression must either occur at the level of BT1927 secretion or trafficking to the outer leaflet of the membrane, its folding to form an effective S-layer or its retention on the cell surface. Interestingly, when we performed additional SEM to attempt to visualize the BT1927 lattice on the *B. thetaiotaomicron* surface (which proved difficult to resolve using the instruments available at the University of Michigan), we noticed a substantial increase in what appeared to be outer membrane vesicles in the supernatant surrounding the *B. thetaiotaomicron* cells (**Figure 5E, F).** *B. thetaiotaomicron* and other *Bacteroides* have been actively investigated in recent years because of their prolific ability to not only produce OMVs but also regulate this process either positively or negatively (45, 46). Quantification of OMVs by a previously published method (47) using the images of fixed and whole-mounted bacterial cultures (with medium supernatant) revealed that growth in GalNAc increases the number of extracellular OMVs ∼25-fold (**Figure 5E, S4F, G**). Thus, while we were unable to visualize the BT1927 lattice with sufficient resolution after growth in fructose and GalNAc, our data suggest that GalNAc induces increased vesiculation by this species which could interfere with the ability of BT1927 to block phage infection.

### The oxidative branch of the pentose phosphate pathway regulates susceptibility to phage

Carbohydrate utilization by *B. thetaiotaomicron* and other *Bacteroides* is a highly regulated process involving multiple layers of regulation. Polysaccharide utilization occurs via expression of substrate-specific PULs and these gene clusters typically exhibit large increases in transcription (50 to >1,000-fold) in response to their target polysaccharide (48). PUL regulation is most commonly mediated by locally acting, positive-acting transcription factors (1). However, global regulatory phenomena—which are still incompletely understood—also exist to control both polysaccharide and monosaccharide utilization in *B. thetaiotaomicron* and at least polysaccharide utilization is subjected to catabolite repression-like effects (*i.e*., some polysaccharides are given high priority while utilization of others is repressed until favored substrates are depleted) (49). Several features involved in global regulation have emerged in recent years, one being a homolog of the *E. coli* catabolite repression protein (CRP) currently referred to as carbohydrate utilization regulator (Cur, previously MalR, *BT4338*). Cur is a global regulator that is required for the normal utilization of multiple poly- and monosaccharides and is thought to work independently of cAMP, which has not been observed in *Bacteroides* (50). Mutants in the oxidative branch of the pentose phosphate pathway (PPP), including 6-phosphogluconate dehydrogenase (BT1222), have been discovered in multiple studies to play a role in polysaccharide or glucosinolate utilization, because these mutants fail to properly express the corresponding PULs or their regulators (50–52). One study found that mutants lacking either functional Cur or the oxidative PPP both fail to fully express a hybrid two component system regulator (Roc) involved in mucin *O*-glycan utilization, implying that these two features may overlap at least partially (50). Interestingly, Roc expression, which is Cur-dependent, is suppressed by glucose and fructose, the two sugars that promote the highest resistance to ARB25.

To determine whether mutants in the PPP or Cur influence ARB25 resistance, we measured growth of four previously identified transposon mutants with disrupted genes in the oxidative branch of the PPP (in the same strain background lacking all 8 capsular polysaccharides used for other experiments in this study). These mutants, which do not have the *BT1927* promoter locked on, exhibited impaired growth relative to the untreated acapsular *B. thetaiotaomicron* strain when infected with ARB25, suggesting a deficiency in deployment of the multiple phase-variable mechanisms that it still retains, including BT1927 (**Figure 6A, Figures S5A-E**). In light of these findings, we evaluated the effects of deleting the *BT1222* gene, encoding the terminal step in the oxidative branch of the PPP, in the BT1927:ON strain. Deletion of *BT1222* resulted in diminished resistance to ARB25 compared to the uninfected control when grown in fructose (**Figure 6B**). Consistent with its apparent increased susceptibility in liquid medium containing fructose, the Δ*BT1222* BT1927:ON strain displayed an increased number of plaques compared to the BT1927:ON parent, albeit less than the parent acapsular strain when both were grown in medium with fructose (**Figure S5F**). Given the increased susceptibility associated with deletion of *BT1222*, we measured *BT1927* transcript levels and confirmed that expression remained unaltered in the Δ*BT1222* mutant (**Figure 6C**). Similar to GalNAC grown cells that showed increased ARB25 susceptibility, we also did not see an appreciable decrease in BT1927 protein by western blot (**Figure S5G**). Finally, using flow cytometry, we observed a decrease in BT1927 surface staining when the Δ*BT1222* BT1927:ON^F^ strain was grown in fructose (**Figure 6D**).

**Figure 6.**
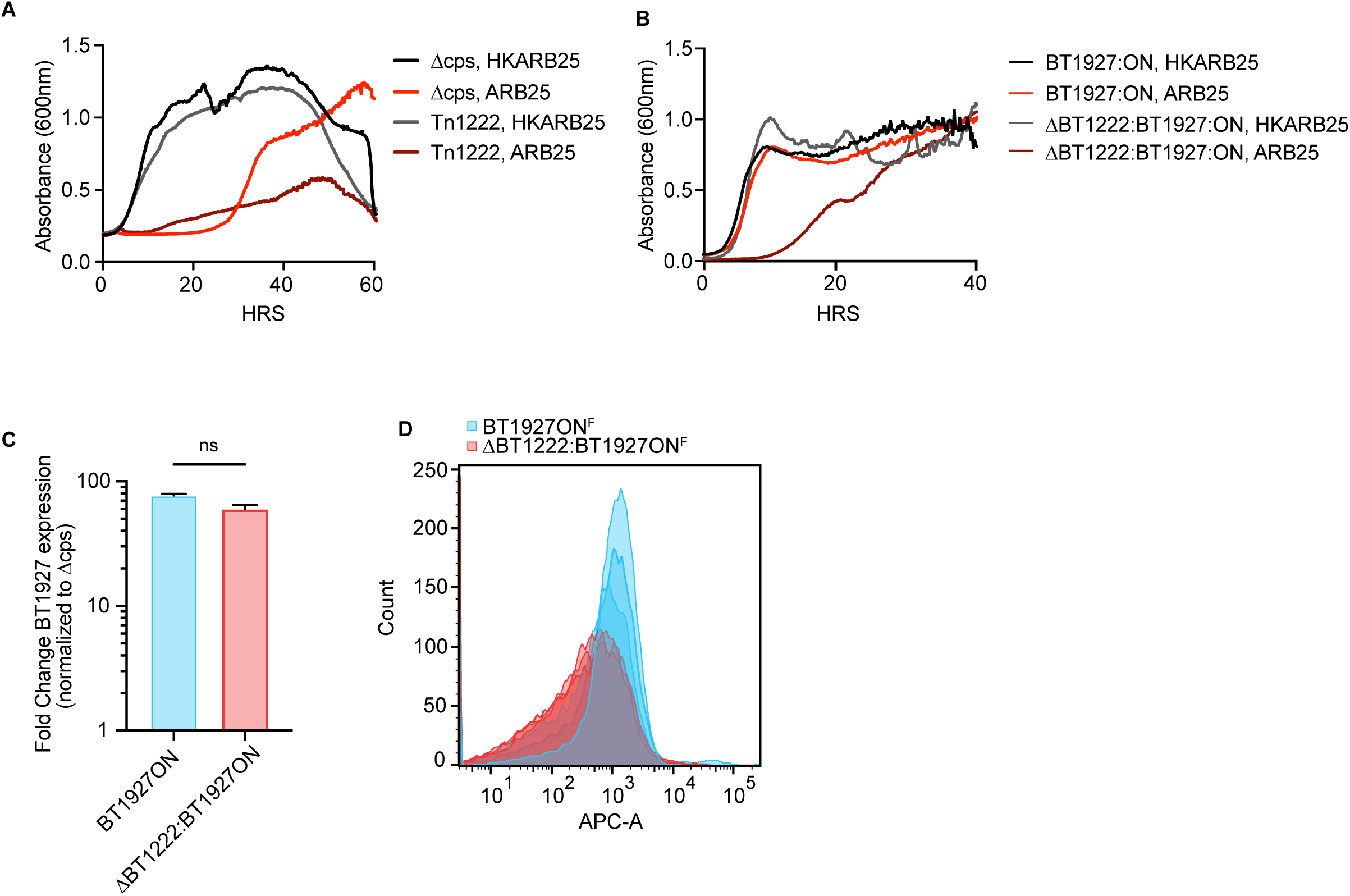
Mutants in the oxidative branch of the pentose phosphate pathway increase susceptibility to ARB25 phage. (**A**) Growth of acapsular *B. thetaiotaomicron* or a transposon mutant in this same strain with disrupted *BT1222* and treated with HK ARB25 (black) or ARB25 (red). (**B**) Growth of the BT1927:ON strain (black) or a *BT1222* deletion (grey) mutant in the BT1927:ON strain background and treated with HK ARB25 or ARB25 (red or dark red, respectively). (**C**) qPCR measurement of *BT1927* transcript in the BT1927:ON strain (light blue) or BT1222 mutant (light red). (**D**) Histograms of epitope-tagged BT1927 (APC-A) for either the BT1927:ON strain (blue) or BT1222 mutant (red) grown in fructose.

We next examined the role of the global regulator Cur, which appears to share some regulation in common with the enzymes in the oxidative branch of the PPP (50). Mutants in which the gene encoding Cur (*BT4338*) was deleted in the Dcps or BT1927:ON backgrounds did not increase susceptibility during growth in fructose as substantially as the PPP mutants, although some increased ARB25 susceptibility was observed (**Figure S5H, I).** Consistent with this, flow cytometry of the epitope-tagged BT1927:ON^F^ strain lacking Cur also displayed similarly intense staining as the parent strain grown in fructose (**Figure S3J**) with an average geometric mean fluorescence intensity of 293±60 (BT1927:ON^F^) and 209±44 (Δ*BT4338* BT1927:ON^F^, P value = 0.32). Taken together, we conclude that the presence of an intact oxidative branch of the PPP is required for fructose- and BT1927-mediated phage resistance, but Cur appears to play a less substantial contribution.

## Discussion

Phages are the most numerous biological entities in many microbial environments, including the human gut (53). The presence of at least some individual phage is stable over time (54), raising the question of how phage can remain present without eradicating their host bacteria or being lost when their host gains resistance. While several mechanisms like lysogeny and pseudolysogeny factor into these relationships, multiple *Bacteroides* species have been shown over the last several decades, using both *in vitro* and *in vivo* experiments, to co-exist with lytic phage in a state where neither completely wins or loses (27, 30). This phenomenon is driven by the existence of phase-variable resistance mechanisms that presumably are selected for in their “on” state when phage are present but revert to the “off” state at a sufficient frequency to continuously generate susceptible prey bacteria to be infected and replenish phage populations. Indeed, in our previous work we showed that ARB25 phage could persist in the colons of *B. thetaiotaomicron* monoassociated mice for over two months, suggesting that reversion of resistant bacteria back to a susceptible state could be sufficient to promote stability of phage populations over longer periods (27). In this study, we bypassed the contributions of phase-variation by locking BT1927 expression in the on state and observed additional, sugar-mediated contributions to *B. thetaiotaomicron* survival, revealing another layer of regulation that influences phage resistance.

The carbohydrate landscape in the mammalian gut is complex and dynamic and also directly influences the metabolism of the bacteria that compose the gut microbiota. Diet is a major external force that shapes the microbiota because dietary fiber—which mostly encompasses plant-derived polysaccharides other than soluble starch—is the major fraction of human foods that reach the colon unabsorbed or undigested by human enzymes (1, 40). *B. thetaiotaomicron* is a nutritional generalist that can utilize a variety of dietary fibers but also has the ability to consume the diverse *O*-glycans attached to secreted gastric and colonic mucin (39, 55). We have previously shown that most dietary fiber polysaccharides that are accessible by *B. thetaiotaomicron* repress expression of this organism’s PULs involved in mucin *O*-glycan degradation through a phenomenon resembling carbon catabolite repression (49). When dietary fiber is withheld from *B. thetaiotaomicron* colonized gnotobiotic mice, there is increased expression of PULs involved in *O*-glycan utilization (41, 56) and measurably increased mucus consumption that manifests as thinner colonic mucus (57). Combined with the results presented here, these observations raise the idea that low dietary fiber could force *B. thetaiotaomicron* into conditions that make it more susceptible to phage predation, such as utilization of *O*-glycans and GalNAc, which both increase its susceptibility to phage like ARB25 (**Figure 3D, E**). This increased susceptibility occurs even though *B. thetaiotaomicron* possesses alternative, mainly phase-variable strategies to resist phage (capsular polysaccharides, putative S-layers, plus others). Indeed, we demonstrate with a broader set of phages that infect *B. thetaiotaomicron,* that GalNAc induces an apparent state of increased susceptibility as judged by reduced growth over time in the presence of several phages (**Figure S2D**), a phenomenon that could be mediated by the increased vesiculation that we note during growth in GalNAc (**Figure 5E, F**). Moreover, we show that increased or decreased susceptibility is entrained by prolonged exposure to different sugars like fructose or GalNAc, revealing that the recent “nutritional history” experienced by *B. thetaiotaomicron* may also play a role. The mechanism(s) through which this nutritional memory occurs remains unknown, although some candidates are noted below, but will be important to investigate and may influence other aspects of *B. thetaiotaomicron* physiology.

Perhaps one of the most unexpected observations in this study, is the detection of what appear to be cell fate decisions in *B. thetaiotaomicron* that are manifest at the single cell level. Evidence to support this comes first from SEM imaging of BT1927:ON and BT1927:OFF strains and the observation that as many as ∼40% of BT1927:ON cells lack the lattice-like surface structure that was previously associated with BT1927 expression and instead resemble the BT1927:OFF strain. Additional evidence to support this idea comes from the observation that some cultures of the BT1927:ON^F^ strain grown in fructose harbor a significant population of cells that resemble the BT1927:OFF by flow cytometry. It has been widely reported that *Bacteroides* species encode a variety of mechanisms to generate phenotypic diversity in the population (27, 58–60). This diversity pre-adapts sub-populations to survive challenges like host innate and adaptive immune responses and phage. In addition to the phase-variable CPS, S-layer, and other functions noted above, which directly impact resistance to phage and immune components, additional phase-variable events include the existence of phase-variable DNA methylation systems (58) that alter a variety of cell features on a global scale, including CPS expression. While these systems have only been investigated in *B. fragilis*, *B. thetaiotaomicron* has two copies of the locus encoding this system (*BT4516-4523* and *BT4535-BT4543*) and likely exhibits similar epigenetic regulation via methylation. In addition, phase-variable gene shufflons that diversify the particular nutrient receptors that are expressed via a single transcription factor exist in *B. thetaiotaomicron* (58). Indeed, we previously observed a pronounced shift in one of these nutrient receptor shufflons *in vitro* after *B. thetaiotaomicron* was exposed to ARB25 and a resistant cell population was allowed to emerge. Such events that directly alter the identities of nutrient receptors on the cell surface could impact phage infection by modifying receptors and co-receptors that mediate adsorption. Although, it is important to note that a test of this hypothesis for ARB25 and the nutrient receptors that we observed to shift upon infection (by deleting the genes involved) did not support a direct role as the ARB25 receptor. Finally, a recent study demonstrated the presence of much more prevalent DNA inversions (372 different “inverton” events) within the *B. thetaiotaomicron* genome and these often occur within coding sequences and are not associated with known recombinases that mediate the inversion (61). While it remains to be investigated, this large amount of genomic variation holds the potential to record aspects of the nutritional memory that we report here, for example, by shifting the genome organization into a different methylation or recombination state that then takes time to shift back when cells encounter another substrate.

It is further surprising that the presence of BT1927 on the cell surface is not regulated at the transcriptional or translational levels, which are more traditional control points to conserve energy. Because cellular levels of BT1927 are very similar during growth in fructose and GalNAc, and only fructose-grown cells are capable of displaying the highest amounts of surface BT1927 staining, this necessitates that GalNAc either reduces the efficiency of BT1927 trafficking to the cell surface or causes it to be released from the surface once it gets there. Indeed, our observation that GalNAc induces increased vesiculation provides evidence that either BT1927 is being lost from the surface, or its protective effects are disrupted by the vesiculation that occurs during growth in GalNAc. These observations also suggests that the mechanism of reduced ARB25 resistance during growth in GalNAc is not identical to the mechanism that sometimes generates the BT1927-off population in cells grown in fructose, or else we would expect the GalNAc population to stain more weakly and identical to the BT1927:OFF^F^ strain.

We do not yet understand the mechanism by which BT1927 protects against phage or, in previous work, against complement-mediated killing. Given the prominent structural organization that is observed on the cell surface when this protein is highly expressed, an obvious hypothesis is that it physically blocks access to either complement deposition and pore formation or phage adsorption. Since the ARB25 surface receptor remains unknown, it also remains unclear how much of the bacterial surface needs to be covered by BT1927 to protect the cell. Indeed, this might not be an all-or-nothing event, but rather, higher percent surface coverage decreases the odds that a phage will access its receptor and infect the cell. Such a phenomenon could explain the higher plaquing rates and clearer plaques we observe with GalNAc-grown cells as well as the “intermediate” growth we often observe in liquid cultures of BT1927:ON grown in GalNAc or conditions containing both GalNAc and fructose that neither achieves full growth or full lysis. In the latter case, growth of protected cells and killing of unprotected cells may balance each other, leading to little positive or negative change in cell numbers measured by absorbance. Future work to identify the ARB25 receptor will open avenues to begin exploring these potential mechanisms in more detail, for example, by increasing or decreasing the amount of ARB25 receptor and seeing how those changes influence killing during growth in fructose or GalNAc.

Our research supports a model of dynamic interactions between phages and intestinal bacteria, which is influenced by the carbohydrates present in the gut. The reduced effectiveness of resistance proteins like BT1927 in the presence of nutrients like GalNAc and *O*-glycans may explain why *B. thetaiotaomicron* possesses so many different strategies (*i.e*., other resistance strategies like capsules may be more effective and therefore relied upon in these growth conditions). Nevertheless, the ability of a phage to persist in the microbiome alongside its prey bacterium like *B. thetaiotaomicron* has implications for what cellular states (*e.g*., which CPS and S-layers are expressed) the resistant bacterial population adopts during selection in the presence of phage. Cell surface features like CPS and S-layers have other interactions with environmental features like the host immune system and fiber polysaccharides (36, 59, 62–64) and, as shown here, are further influenced by the carbohydrates *B. thetaiotaomicron* utilizes. With this complex set of interactions between phage, nutrients, bacteria and host in mind, it is likely that the variable presence of phage within the microbiomes of individuals has broader impacts on holobiont physiology that extend beyond just simple survival of the bacteria that are targeted by phage.

## Materials and Methods

### Bacterial growth curves with phage

Table 1 provides a list of strains used in this study. All *B. thetaiotaomicron* cultures were grown in an anaerobic chamber (Coy Labs; 85% N2, 10% H2, 5% CO2) at 37°C. In general, growth experiments were initiated by streaking freezer stocks (cryopreserved in 25% final w/v glycerol) on *Bacteroides* phage recovery medium (BPRM) agar with glucose as the main carbon source (66). High titer phage stocks were propagated as previously described (27, 67). Briefly, the acapsular *B. thetaiotaomicron* strain was incubated with 10ul of high titer phage for 20 min and plated using a soft agar overlay, flooded after overnight incubation using sterile phage buffer (100 mM NaCl, 8 mM MgSO4, 50 mM Tris pH 7.5, and 0.002% (w/v) gelatin), filter sterilized, and the resulting phage titer determined using the acapsular strain as the host.

**Table 1:** Bacterial strains and plasmids used in this study.

| Table 1: Bacterial strains and plasmids used in this study |  |  |  |
| --- | --- | --- | --- |
| Strain | Genotype | Features | Reference |
| ***Bacteroides thetaiotaomicron* VPI-5482 and derivatives** |  |  |  |
| acapsular | *tdk Δcps1-8* | Strain lacking all CPS synthesis loci | (65) |
| acapsular S-layer ON | *tdk Δcps1-8*<br>BT1927:ON | Strain lacking all CPS synthesis loci, S-layer promoter locked in the ON orientation | (27) |
| acapsular S-layer OFF | *tdk Δcps1-8*<br>BT1927:OFF | Strain lacking all CPS synthesis loci, S-layer promoter locked in the OFF orientation | (27) |
| acapsular S-layer ON FLAG | *tdk Δcps1-8*<br>BT1927:ON FLAG | Strain lacking all CPS synthesis loci, S-layer promoter locked in the ON orientation with the C-terminal FLAG epitope | This study |
| acapsular S-layer OFF FLAG | *tdk Δcps1-8*<br>BT1927:OFF FLAG | Strain lacking all CPS synthesis loci, S-layer promoter locked in the OFF orientation with the C-terminal FLAG epitope | This study |
| ΔBT1222 acapsular | *tdk Δcps1-8*<br>BT1927:ON | Strain lacking all CPS synthesis loci, gene deletion of *BT1222* | This study |
| ΔBT1222 acapsular S-layer ON | *tdk Δcps1-8*<br>BT1927:ON ΔBT1222 | Strain lacking all CPS synthesis loci, S-layer promoter locked in the ON orientation, gene deletion of *BT1222* | This study |
| ΔBT1222 acapsular S-layer ON FLAG | *tdk Δcps1-8*<br>BT1927:ON ΔBT1222 FLAG | Strain lacking all CPS synthesis loci, S-layer promoter locked in the ON orientation with the C-terminal FLAG epitope and gene deletion of *BT1222* | This study |
| ΔBT4338 acapsular | *tdk Δcps1-8* ΔBT4338 | Strain lacking all CPS synthesis loci, gene deletion of *BT1222* | This study |
| $\Delta BT4338$<br>acapsular S-layer ON | *tdk* $\Delta cps1-8$<br>BT1927:ON $\Delta BT4338$ | Strain lacking all CPS synthesis loci, S-layer promoter locked in the ON orientation, gene deletion of *BT1222* | This study |
| $\Delta BT4338$<br>acapsular S-layer ON FLAG | *tdk* $\Delta cps1-8$<br>BT1927:ON $\Delta BT4338$<br>FLAG | Strain lacking all CPS synthesis loci, S-layer promoter locked in the ON orientation with the C-terminal FLAG epitope and gene deletion of *BT1222* | This study |
| **Plasmids in *Escherichia coli* S17-1 λpir, used for generation of *B. thetaiotaomicron* gene deletions** |  |  |  |
| pExchange<br>BT1927-ON |  | Lock on BT1927 promoter | (36) |
| pExchange<br>BT1927-OFF |  | Lock off BT1927 promoter | (36) |
| pExchange<br>BT1927- ON<br>FLAG |  | Lock on BT1927 promoter with C-terminal FLAG epitope | (36) |
| pExchange<br>BT1927- OFF<br>FLAG |  | Lock off BT1927 promoter with C-terminal FLAG epitope | (36) |
| pExchange<br>$\Delta BT1222$ | | *BT1222* deletion construct | This study |
| pExchange<br>$\Delta BT4338$ | | *BT14338* deletion construct | This study |

Individual colonies of each strain were picked from BPRM agar after 3 days and grown overnight in liquid BPRM medium with glucose as the main carbon source, unless otherwise indicated. For growth experiments, an aliquot of each overnight culture was centrifuged and washed twice with a 2X concentrated BPRM without any glucose or other carbon source (BPRM no carbon). The washed cells were resuspended in BPRM no carbon and 50ul was added to a 96-well microtiter plate wells containing 10mg/ml of carbohydrate as indicated in each growth curve, for a final concentration of 5mg/ml. 10ul of 5×10^6^ pfu/ml stock of either heat-killed (95°C for 30 min) or viable phage was added to each well. Microtiter plates were covered with a gas-permeable, optically clear membrane (Diversified Biotech), and optical density measured at 600nm (OD_600_) for 24-96 hours using a BioTek Synergy HT plate-reader connected to a BioStack automated plate stacking device.

### Construction of *B. thetaiotaomicron* mutants

All of the *B. thetaiotaomicron* mutants were created in the acapsular *B. thetaiotaomicron (*Δcps Δtdk) strain, a mutant for thymidine kinase used to generate allelic exchange mutants (68) and lacks all capsule. *B. thetaiotaomicron* Δcps BT1927:ON and Δcps BT1927:OFF mutants were created from the previously constructed BT1927-ON and BT1927-OFF strains (36) and the corresponding epitope FLAG tagged constructs for each. The *BT1222* and *BT4338* gene deletion mutants were generated by homologous regions for allelic exchange (68) with primers indicated in Table 2. All mutants were confirmed by PCR and Sanger sequencing.

**Table 2.**
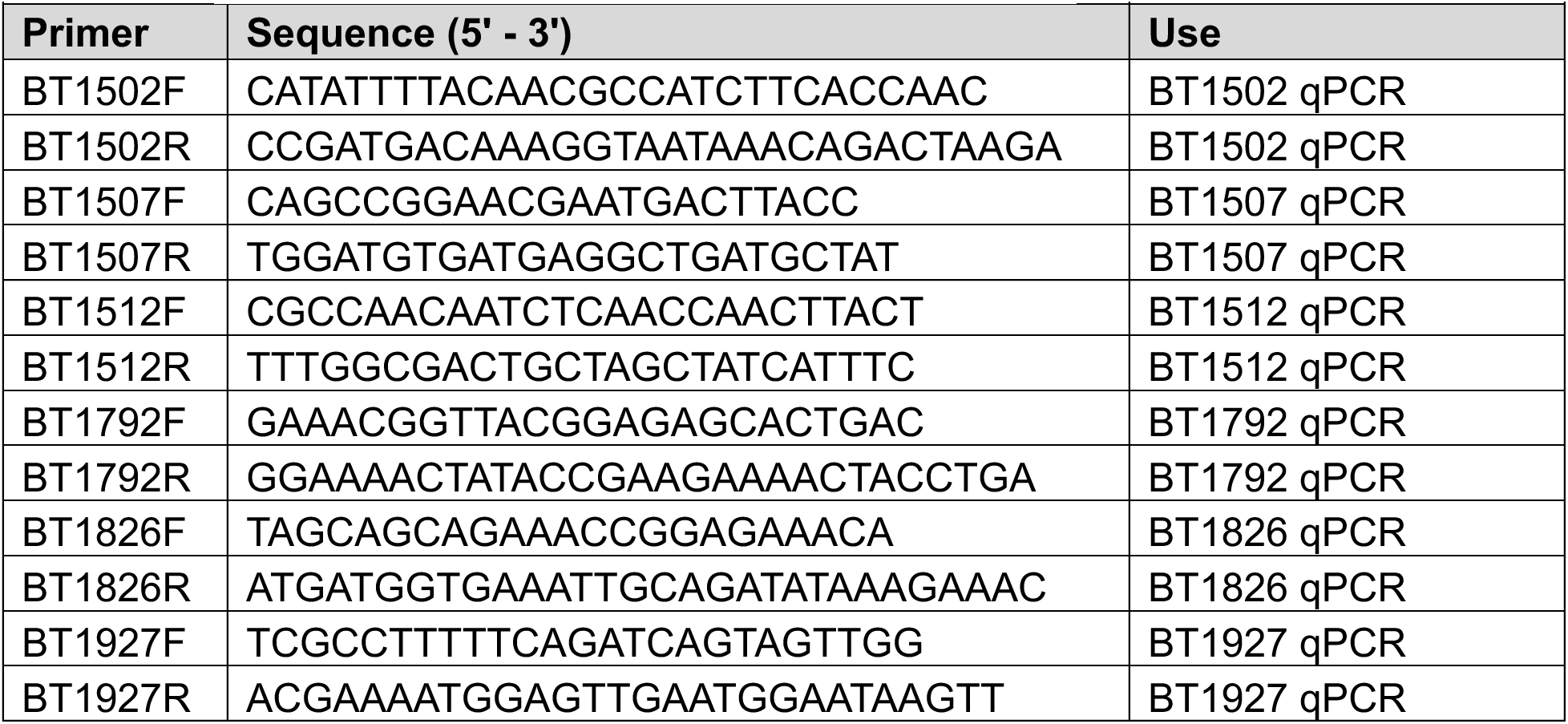
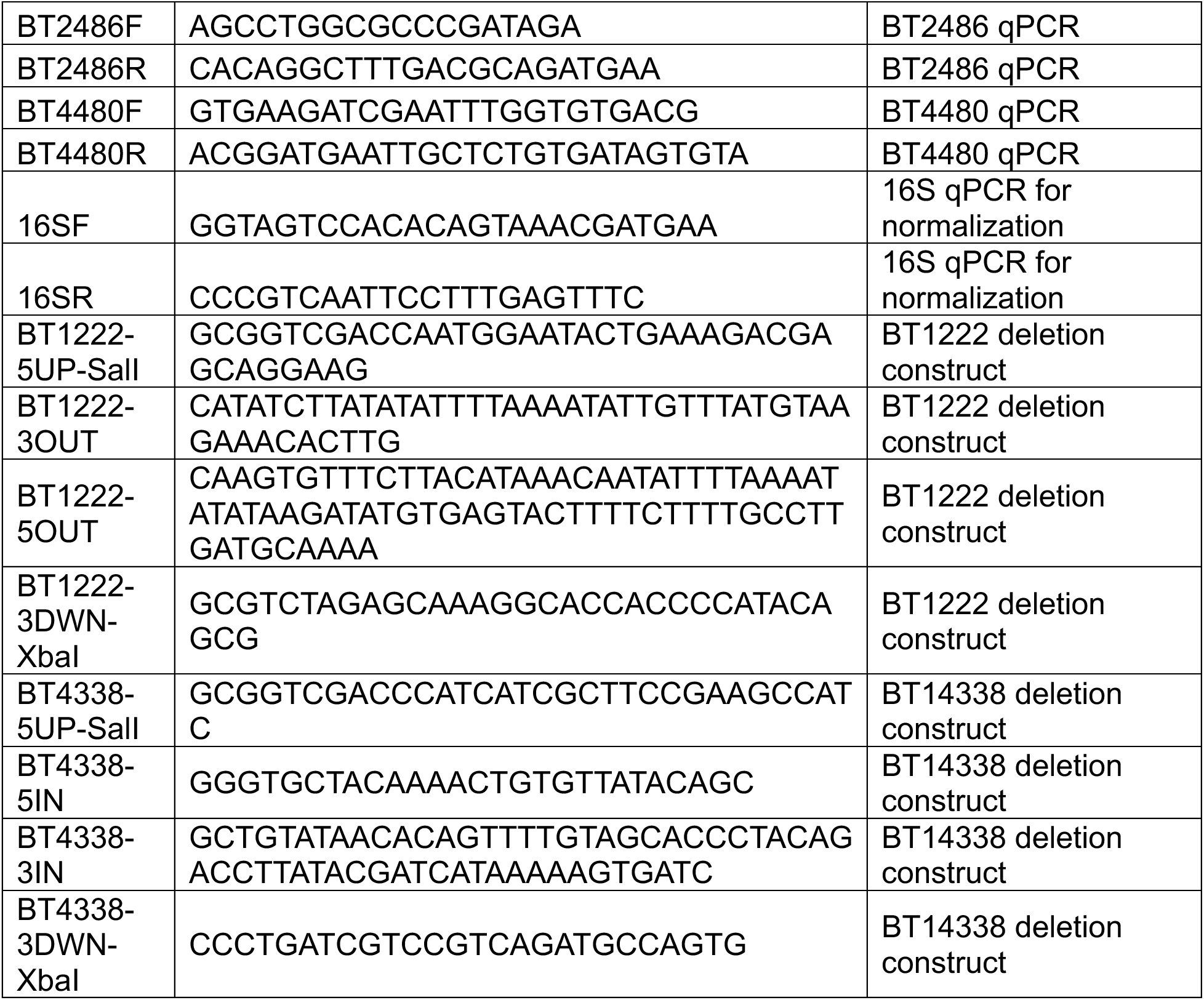
Primers used in this study.

### Plaque Assay

Isolated colonies of the BT1927:ON strain were cultured overnight in liquid BPRM media and added to soft agar overlay as previously described (27) containing either glucose, fructose or GalNAc as indicated. High titer ARB25 stocks were diluted 1:10 in series and 1ul was added to the soft agar containing the BT1927:ON or Δcps strain, incubated overnight anaerobically, and quantified. To document plaque morphology, images were captured using the GelDoc Go Gel Imaging System (Bio-Rad) using a 0.8 sec exposure.

### Complement Survival Assay

*B. thetaiotaomicron* strains were cultured overnight as described above from isolated colonies in liquid BPRM media containing glucose. Cultures were prepared as previously described (36). Briefly, Cells were centrifuged and washed once with PBS and resuspended at 5 × 10^6^ cells/ml in PBS supplemented with 0.5 mM MgCl2 and 1 mM CaCl2. Either 20% Human Serum or heat-inactivated human serum (56°C for 30 min) was incubated with cells anaerobically at 37°C for 1 hour. Cells were diluted in series 1:10 and plated on BHI plates containing 10% horse blood anaerobically for 2 days. The percent survival of each strain was calculated by dividing the average number of colonies from human serum vs heat-inactivated serum plates.

### RNA isolation and quantitative PCR (qPCR)

To measure BT1927 and other gene transcript levels, cultures were grown as described above in BPRM containing the different sugars indicated until they reached OD_600_ between 0.5-0.6. Bacteria were collected by centrifugation (7,700xg for 2.5 min), medium supernatants were removed, and pellets were resuspended in 500uL of RNAprotect (Qiagen). RNA was extracted using RNeasy kit (Qiagen) and treated with DNase I (NEB). cDNA was prepared using 1ug RNA via Superscript III Reverse Transcriptase according to the manufacturer’s instructions. qPCR analyses were performed using a BioRad thermocycler and homemade qPCR Master Mix containing SYBR green DNA dye (27) containing 500nM of BT1927 specific primers or 65nM 16S rRNA primers (indicated in table 2) and 10ng cDNA. Amplification was performed for 40 cycles of 95°C for 3 seconds, 55°C for 20 seconds, and 72°C for 8 seconds. Normalization was performed using the 16S rRNA amplification and gene expression between conditions was calculated using the DD cycle threshold (DDCt) method (69) using the Δ*cps* strain, which expresses little *BT1927* in the absence of phage, as a reference.

### Flow cytometry

The *B. thetaiotaomicron* strain containing a FLAG epitope-tagged BT1927 locked-on strain was previously constructed (36) using the FLAG epitope tag to BT1927 C-terminus. Various strains containing this tagged allele were cultured as described above with the addition of indicated carbohydrates and grown to OD_600_ between 0.5-0.6. One milliliter of culture was centrifuged for 2 min in a microcentrifuge at 6,000 x g, washed twice with PBS, and resuspended to approximately 5×10^6^ cells/ml in Bacteria Staining buffer (BSB; 1% bovine serum albumin– 0.025% sodium azide–PBS, 0.5% formaldehyde) and resuspended in BSB containing anti-FLAG Alexa Fluor 647-conjugated antibody (1:25, Cell Signaling) and incubated for 1h at 37°C. Cells were washed twice with BSB and were counterstained with SYTO BC Bacteria Stain (Molecular Probes) and analyzed on an LSR Fortessa flow cytometer (BD). To minimize carryover of cells, 3 blank samples of PBS were run between samples. Forward scatter (FSC) and side scatter (SSC) data were set to logarithmic scale. After data acquisition, *B. thetaiotaomicron* population was visualized on an allophycocyanin (APC-A)-histogram plot using FlowJo software.

### Immunoblotting

*B. thetaiotaomicron* strains containing the FLAG epitope-tagged BT1927 allele (36) were cultured as described above with the addition of indicated carbohydrates and grown to OD_600_ between 0.5-0.6. Samples were diluted 1:4 in Laemmli sample buffer (Bio-Rad), heated for 10 min at 98°C and were run on a 4-12% Bis-Tris NuPAGE gel (Invitrogen) and transferred onto a polyvinylidene difluoride membrane (0.2 µm) for Western blot analysis. Membranes were blocked using Tris-buffered saline tween (TBST)-based blocking solution (1M Tris, 5M NaCl, Tween 20) containing 5% milk, stained with the anti-FLAG rabbit polyclonal primary antibody (1:1000, Cell Signaling), washed twice with TBST, and goat polyclonal anti-rabbit (ThermoFisher) conjugated Alexa Fluor 488 secondary antibody (1:1000). Imaging was performed using an Odyssey CLx scanner (LI-COR).

### Proteomics

The BT1927 locked-on strain was grown in BPRM containing either fructose or GalNAc to OD_600_ between 0.5-0.6., cells were centrifuged (7,700xg for 2.5 min), washed in PBS and frozen at −80°C. Cell pellets were resuspended in 500µl lysis buffer (4% SDS, 50mM Tris, 10mM DTT, pH8.5), heated to 95°C for 10 min, cooled to room temperature, and centrifuged for 2 min at 4,000 x g to remove cell debris and unlyzed cells. Ice-cold acetone was added at 4:1 ratio and precipitated overnight at −20°C. Proteins were centrifuged for 10 min at 4,000 x g, and the resulting pellets were washed in 80% acetone before centrifuging at 17,000 x g for 10 min at 4°C. Supernatants were removed, and the protein pellet allowed to air dry before resuspension in 75µl of proteomics buffer (8M Urea, 50mM HEPES, pH8). Cell lystes were run on a 4-12% Bis-Tris NuPAGE gel (Invitrogen) and subjected to Coomassie staining (Abcam). Proteomics analysis was performed at the University of Michigan Proteomics Resource Facility (PRF) using high-resolution LC-MS/MS, following a protocol refined by PRF as previously described (70). The updated protocol included the following changes: cysteines were alkylated with 65 mM 2-chloroacetamide; trypsin digestion occurred after urea was diluted to <1.2 M, utilizing 0.5 µg of trypsin (Promega); and processed peptides were reconstituted in 20 µL of Buffer A (100 mM NaOH). Peptides were separated by liquid chromatography over 90 minutes in 2%–25% Buffer B (100 mM NaOH and 500 mM sodium acetate) in 45 minutes, 25%–40% in 5 minutes, and 40%– 90% in 5 minutes, followed by holding at 90% buffer B for 5 minutes and equilibrating with buffer A for 30 minutes). Proteins were identified by comparing the resulting tandem mass spectrometry data against a database of all protein sequences in *B. thetaiotaomicron* strain VPI-5482 (71). The parameters used were consistent with those stated previously, except for a fragment tolerance of 0.1 kDa. The relative abundance of each detected protein was determined based on MS1 abundances. Raw data is available via the Proteomics Identifications Database (PRIDE) # PXD063111.

### Scanning Electron Microscopy

For SEM imaging performed in Tuebingen, the BT1927:ON and BT1927:OFF strains were grown on BHIS plates with 10% defibrinated sheep blood and grown anaerobically for 4 days at 37°C. Individual colonies were picked into liquid BHIS medium and grown for 15 hours at 37°C prior to fixation in 2.5% glutaraldehyde/4% formaldehyde in PBS for 2 hours at room temperature. Sample processing and imaging was performed by the Electron Microscopy Core at the Max Planck Institute for Biology in Tübingen, Germany. Cells were mounted on poly-L-lysine-coated cover slips and post-fixed with 1% osmium tetroxide for 45 minutes on ice. Subsequently, samples were dehydrated in a graded ethanol series followed by critical point drying (CPD300, Leica Microsystems) with CO_2_. Finally, the cells were sputter-coated with a 3 nm thick layer of platinum (CCU-010, Safematic) and examined with a field emission scanning electron microscope (Regulus 8230, Hitachi High Technologies) at an accelerating voltage of 3 kV.

For Cell and OMV imaging at the University of Michigan, 3-day aged colonies of the BT1927 locked-on strain were cultured in BPRM glucose for 20 hours and diluted 1:100 in BPRM containing either fructose or GalNAc to OD_600_ between 0.5-0.6, fixed in 2.5% glutaraldehyde/4% formaldehyde in PBS for 2 hours at room temperature, mounted on poly-L-lysine-coated cover slips, and post-fixed with 1% osmium tetroxide for 45 minutes on ice. Samples were dehydrated in a graded ethanol series followed by critical point drying (CPD300, Leica Microsystems) with CO_2_ and sputter coated 3 nm thick layer of gold and examined with a field emission scanning electron microscope (Thermo Fisher Nova 200 Nanolab SEM/FIB) at an accelerating voltage of 3 kV. OMV particles were quantified by normalized image size, creation of 10×10mm boxes almost devoid of cells, images were blinded and OMV particles were quantified. OMV particle count was divided by the total area to account for any differences in the number of boxes per image.

## Acknowledgments

We acknowledge support from the University of Michigan Biomedical Research Core Facilities (Proteomics Resource Facility and Microscopy core), which is supported by a National Institutes of Health (NIH) Center Grant (University of Michigan Center for Gastrointestinal Research, P30 DK034933). J.J.F was supported by funds from the NIH T32 Genetics Training Program (T32GM007544) and the Howard Hughes Medical Institute Gilliam Fellowship (GT15133) for which we are extremely grateful. We acknowledge the Electron Microscopy Core Facility at the Max Planck Institute for Biology Tuebingen, especially Jürgen Berger and Katharina Hipp, for their expertise and support from the Max Planck Society to SLH and REL. We acknowledge the financial support of the University of Michigan College of Engineering, NSF grant (DMR-0320740), and technical support from the Michigan Center for Materials Characterization.

---

**Figure S1.**
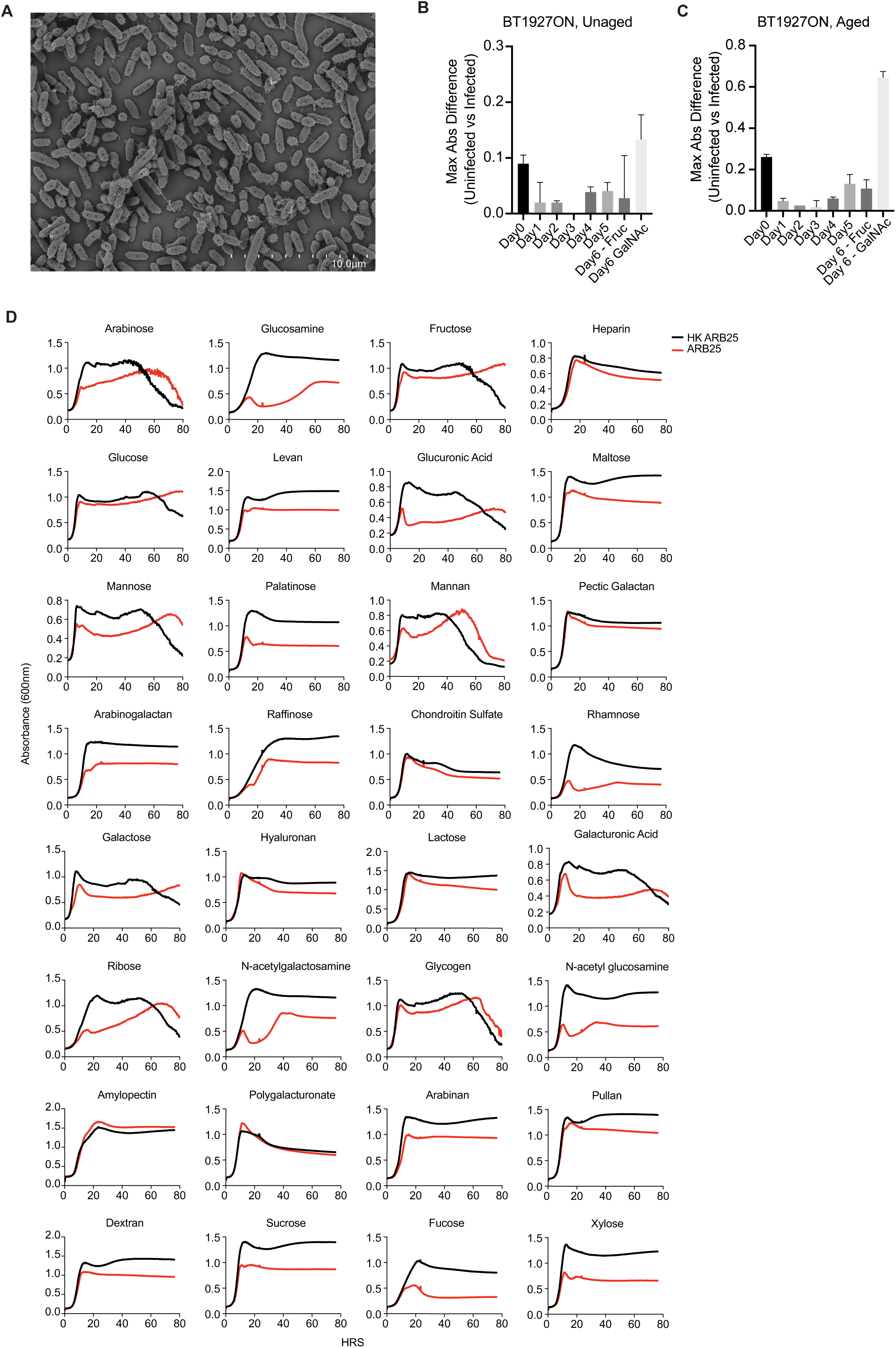
(**A**) SEM imaging of the BT1927:ON strain with cells that exhibit BT1927 associated crystalline coating over most of their surface (pink dot, green circle) and what appear to be uncoated cells (green circle only). (**B**) Maximum absorbance (abs) differences of unaged BT1927:ON after 5-days of passage in glucose and subsequent passage into fructose or GalNAc. (**C**) Maximum absorbance (abs) differences of aged BT1927:ON after 5-days of passage in glucose and subsequent passage into fructose or GalNAc. (**D**) Growth of the BT1927:ON strain in BPRM medium containing the carbohydrates listed above each graph as the sole carbohydrate source (5mg/ml, n= 3 per condition).

**Figure S2.**
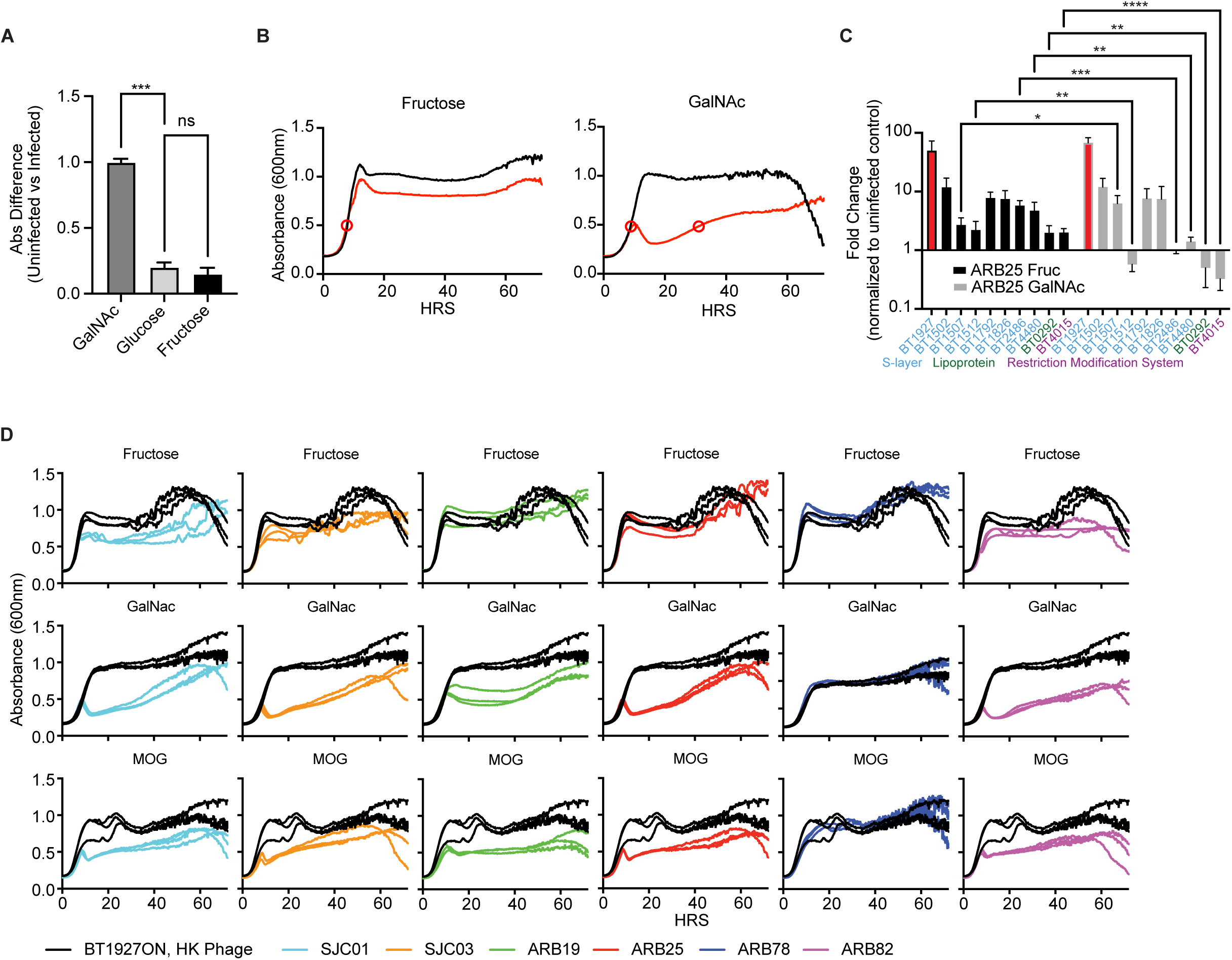
(**A**) Absorbance difference between ARB25 infected and HK control for cultures grown in GalNAc, glucose, or fructose (One-way ANOVA, **** P<0.0001, ***P=0.0002, **P<0.007, *P<0.05). (**B**) Cultures of the BT1927:ON strain grown in fructose or GalNAc for RNA collection with harvest points indicated by red circles in the infected and HK control for each curve. (**C**) Transcript expression measurements for each phase-variable S-layer or other system as measured by qPCR in fructose and GalNAc. (**D**) BT1927:ON cultures treated with HK *B. thetaiotaomicron* phage (black) or challenged with the 6 indicated *B. thetaiotaomicron* phages in BPRM containing fructose (top) GalNAc (middle) or mucin *O*-glycans (MOG; Bottom), n = 3 per condition.

**Figure S3.**
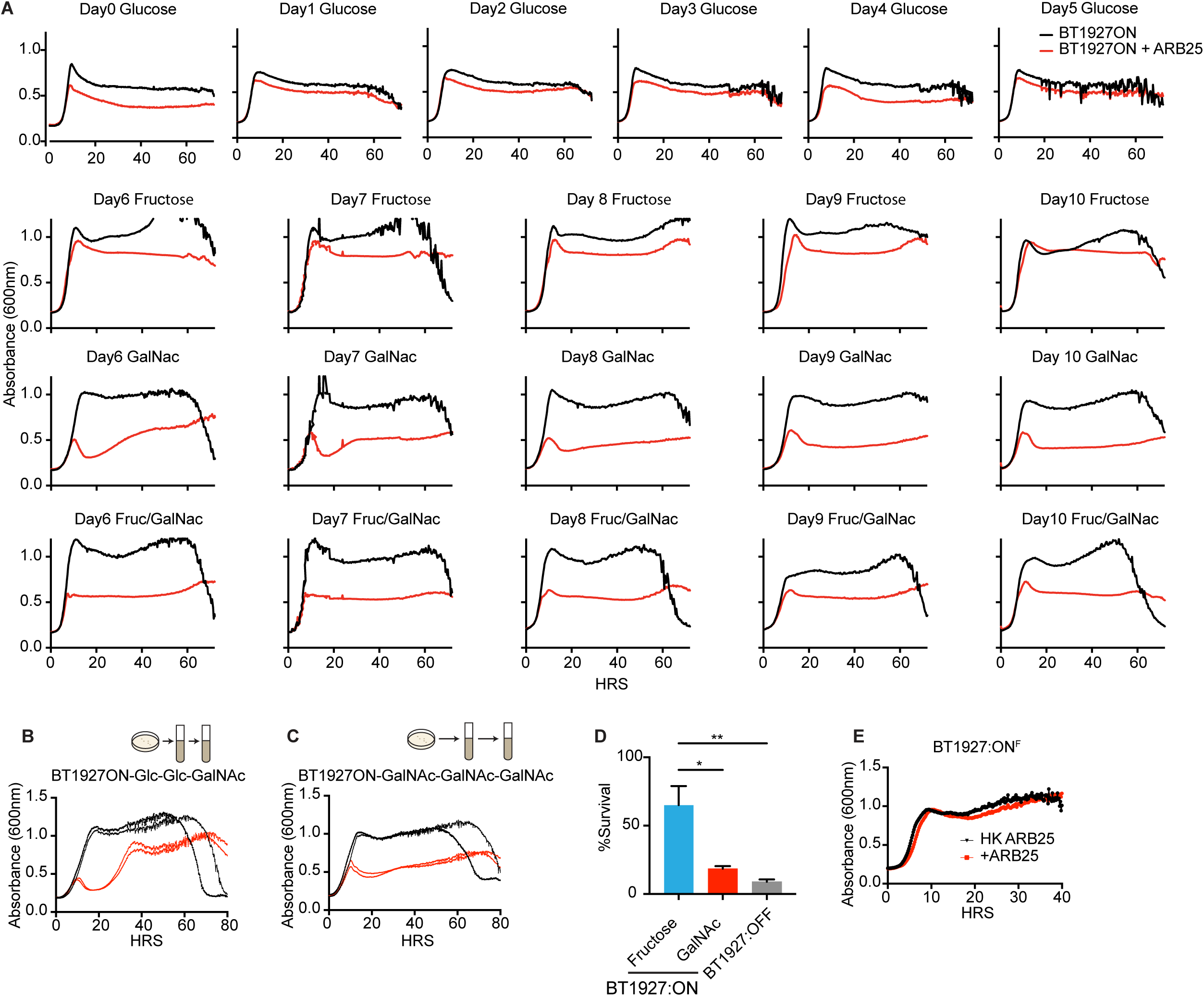
(**A**) Growth of daily passaged BT1927:ON cultures in BPRM medium containing glucose (5 days) and subsequent growth after passage into fructose, GalNAc, or 1:1 mixture of fructose and GalNAc (at 5mg/ml each) as indicated, n = 3 per condition. (**B**) ARB25 infection kinetics for BT1927:ON cultures grown on a solid medium containing glucose, grown in liquid culture containing glucose and sub-cultured again into liquid media containing GalNAc (as indicated above title) and challenged with ARB25 phage. (**C**) ARB25 infection kinetics for BT1927:ON cultures grown on a solid medium containing GalNAc, grown in liquid culture containing GalNAc and sub-cultured again into liquid media containing GalNAc (as indicated above title) and challenged with ARB25 phage. (**D**) Percent survival of BT1927:ON grown in either fructose (blue) or GalNAc (red) or the BT1927:OFF strain grown in fructose (grey) challenged with pooled human complement and normalized to the no complement control. (**E**) Growth of the epitope tagged BT1927:ON^F^ strain treated with HKARB25 (black) or ARB25 (red) demonstrating the FLAG tag does not inhibit BT1927-mediated resistance.

**Figure S4.**
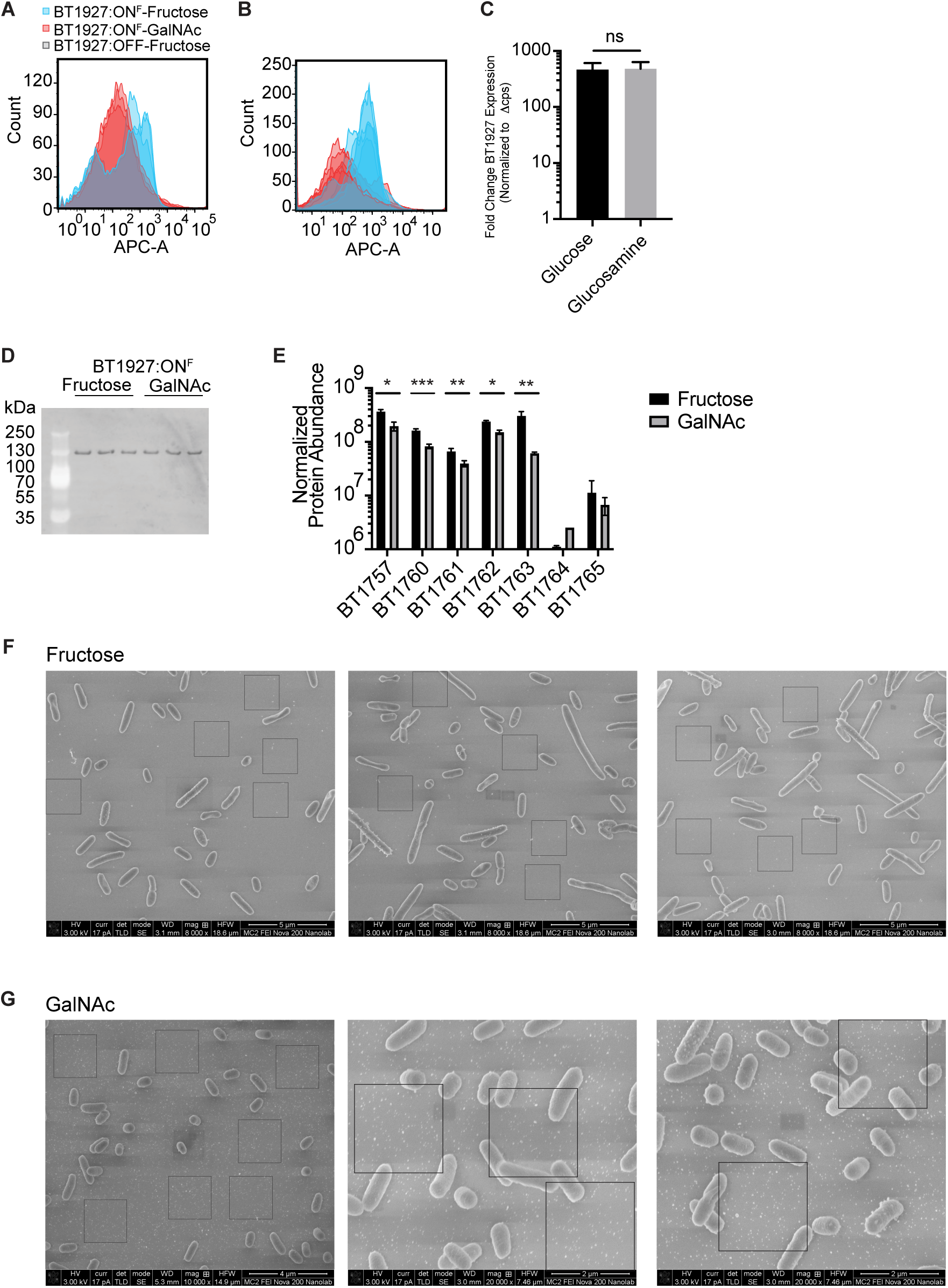
(**A**) Flow cytometry histogram for the BT1927:ON^F^ strain grown in BPRM medium containing fructose or GalNAc from an experiment in which all 3 replicates showed a BT1927 low staining population. (**B**) Flow cytometry histogram for the BT1927:ON^F^ strain grown in BPRM medium containing fructose or GalNAc from an experiment in which replicates were grown in different anaerobic chamber and one showed a BT1927 low staining population. (**C**) *BT1927* transcript expression in the BT1927:ON strain grown in glucose (black) or glucosamine (grey) at mid-log phase. (**D**) Western blot of whole cells of the BT1927:ON^F^ strain grown in either fructose or GalNAc and stained for BT1927-FLAG. (**E**) Normalized protein abundance for products of the levan associated PUL that is known to be induced by fructose (two-tailed t-test, P *<0.001, **<0.05, *** <0.005). (**F**) SEM images of the BT1927:ON strain grown in fructose from which OMV measurements were made. (**G**) SEM images of the BT1927:ON strain grown in GalNAc from which OMV measurements were made. In F and G, boxes indicate normalized area used for blinded counting of OMV particles.

**Figure S5.**
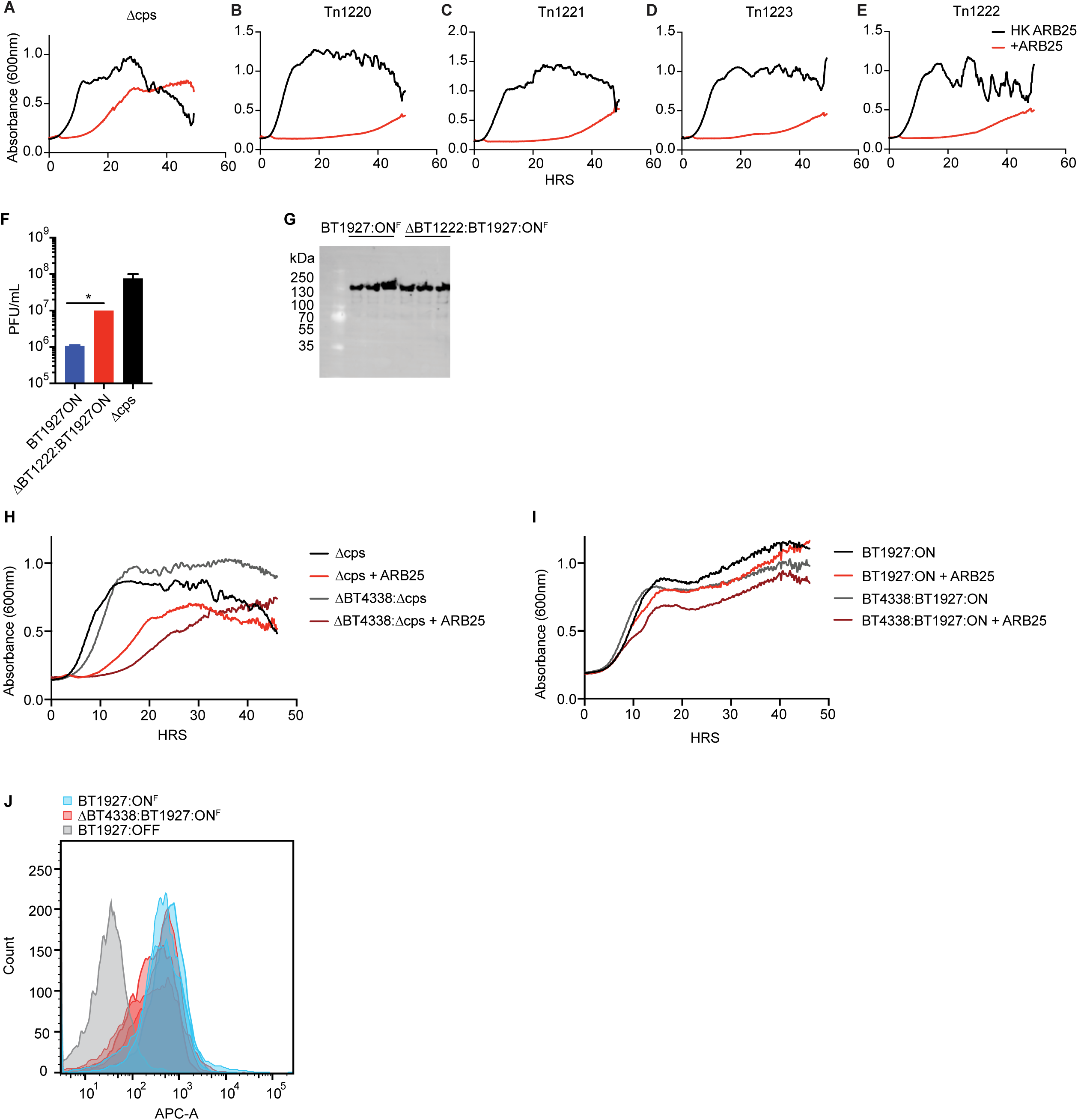
(**A-E**) Growth of the acapsular *B. thetaiotaomicron* strain (A) and transposon mutants in the acpasular *B. thetaiotaomicron* with insertions in genes encoding the 3 steps of the oxidative branch of the pentose phosphate pathway and a linked genes encoding a putative carboxylate transporter. In each panel, cultures were treated with HK ARB25 (black) or ARB25 (red). (**F**) Quantification of plaques formed on the BT1927:ON strain (blue), a *BT1222* deletion in the 1927:ON strain background (red) or the susceptible acapsular strain (black). (**G**) Western blot of the BT1927:ON^F^ or Δ*BT1222* mutant in the BT1927:ON^F^ strain as indicated above each column. (**H**) Growth of the acapsular *B. thetaiotaomicron* or Δ*BT4338* deletion mutant strain in the same acapsular background treated with HK ARB25 or ARB25 (red or dark red, respectively). (**I**) Growth of the BT1927:ON (black) or corresponding Δ*BT4338* deletion mutant in the same background (grey) in the BT1927:ON strain treated with HK ARB25 or ARB25 (red or dark red, respectively). (**J**) Histogram of staining for the epitope-tagged BT1927 (APC-A) in either the BT1927:ON strain (blue) or a Δ*BT4338* mutant in this same background (red), grown in fructose.

